# Physiological noise modeling in fMRI based on the pulsatile component of photoplethysmograph

**DOI:** 10.1101/2020.06.01.128306

**Authors:** Michalis Kassinopoulos, Georgios D. Mitsis

**Author notes:** **Corresponding author:** Michalis Kassinopoulos, Graduate Program in Biological and Biomedical Engineering, McGill University, 350 McConnell Engineering Building, 3480 University Street, Montreal, QC, H3A 0E9, Canada.

## Abstract

The blood oxygenation level-dependent (BOLD) contrast mechanism allows the noninvasive monitoring of changes in deoxyhemoglobin content. As such, it is commonly used in functional magnetic resonance imaging (fMRI) to study brain activity since levels of deoxyhemoglobin are indirectly related to local neuronal activity through neurovascular coupling mechanisms. However, the BOLD signal is severely affected by physiological processes as well as motion. Due to this, several noise correction techniques have been developed to correct for the associated confounds. The present study focuses on cardiac pulsatility fMRI confounds, aiming to refine model-based techniques that utilize the photoplethysmograph (PPG) signal. Specifically, we propose a new technique based on convolution filtering, termed cardiac pulsatility model (CPM) and compare its performance with the cardiac-related RETROICOR (Card-RETROICOR), which is a technique commonly used to model fMRI fluctuations due to cardiac pulsatility. Further, we investigate whether variations in the amplitude of the PPG pulses (PPG-Amp) covary with variations in amplitude of pulse-related fMRI fluctuations, as well as with the systemic low frequency oscillations (SLFOs) component of the fMRI global signal (GS – defined as the mean signal across all gray matter voxels). Capitalizing on 3T fMRI data from the Human Connectome Project, CPM was found to explain a significantly larger fraction of the fMRI signal variance compared to Card-RETROICOR, particularly for subjects with larger heart rate variability during the scan. The amplitude of the fMRI pulse-related fluctuations did not covary with PPG-Amp; however, PPG-Amp explained significant variance in the GS that was not attributed to variations in heart rate or breathing patterns. Our results suggest that the proposed approach can model high-frequency fluctuations due to pulsation as well as low-frequency physiological fluctuations more accurately compared to model-based techniques commonly employed in fMRI studies.

## 1. Introduction

Functional magnetic resonance imaging (fMRI) is a powerful neuroimaging modality that provides measurements of brain activity with great spatial coverage and resolution (Bandettini et al., 1992; Kwong et al., 1992; Ogawa et al., 1990). It has been extensively used in behavioral experiments to study brain function associated to a specific task or condition but also during resting conditions to examine the intrinsic brain functional architecture (Biswal et al., 1995; Van Dijk et al., 2010). The majority of fMRI experiments are based on the blood oxygen level dependent (BOLD) contrast that can detect changes in blood oxygenation, and specifically changes in concentration of deoxygenated hemoglobin (dHb). The main principle exploited in BOLD fMRI is that neuronal activity triggers changes in local cerebral blood flow (CBF) which, in turn, affects the concentration of dHb (Iadecola, 2017; Kisler et al., 2017). As such, due to its sensitivity to levels of dHb, the BOLD signal provides an indirect measure of the underlying neuronal activity. However, the BOLD signal is influenced by physiological-related fluctuations, as well as fluctuations due to motion which, if not properly accounted for, can severely diminish the ability to detect neural-induced signals or lead to biased connectivity profiles (Birn, 2012; Chang and Glover, 2009a; Glasser et al., 2018; Power et al., 2015; Savva et al., 2020; Xifra-Porxas et al., 2020).

The effects of physiological confounds may have important implications for task-based fMRI, as a variety of tasks can lead to task-correlated breathing variations, which in turn give rise to BOLD fluctuations that can be mistaken for neuronally-driven changes (Birn et al., 2009; Glasser et al., 2018). In addition, the effects of physiological confounds are of particular concern for studies investigating the neural correlates of the autonomic nervous system. For instance, a common approach for studying brain-heart interactions is by examining the relationship of variations in functional connectivity with changes in heart rate variability (Chang et al., 2016, 2013; Mulcahy et al., 2019). However, systemic hemodynamic fluctuations induced by changes in heart rate (HR), if not removed from the data, can lead to artificial connectivity with potential implications for the interpretation of the findings.

The fMRI confounds induced by physiological processes and motion fall into two categories: (1) purely physiological blood-borne signals also known as systemic low-frequency oscillations (SLFOs), and (2) acquisition artifacts. Blood-borne signals are signals driven by changes in the levels of dHb in the sample being imaged which in principle can be influenced by several physiological factors. Experimentally, it has been shown that variations in HR (Shmueli et al., 2007), levels of carbon dioxide (CO_2_; Prokopiou et al., 2019; Wise et al., 2004), breathing patterns (Birn et al., 2006), as well as arterial blood pressure (Whittaker et al., 2019) give rise to low-frequency (∼0.1 Hz) fluctuations in fMRI presumably due to their effects on the levels of dHb (Caballero-Gaudes and Reynolds, 2017; Liu, 2016; Murphy et al., 2013). On the other hand, acquisition artifacts are caused by any kind of motion that forces the imaged sample to move in space or perturbs the magnetic field, as these movements have a direct impact on the acquisition process (Caballero-Gaudes and Reynolds, 2017; Liu, 2016; Murphy et al., 2013). Acquisition artifacts may be related to bulk head motion and breathing-related chest expansion (Power et al., 2015) but also to cardiac contractions through vessel expansion in the brain vasculature and its associated tissue movement (Dagli et al., 1999).

To account for the effects of physiological processes and motion, several data-driven techniques have been proposed (Caballero-Gaudes and Reynolds, 2017). An important class of data-driven techniques involves the decomposition of the fMRI data into components using either principal or independent component analysis (Behzadi et al., 2007; Pruim et al., 2015; Salimi-Khorshidi et al., 2014) and the removal of components that are likely due to physiological fluctuations or acquisition artifacts prior to further analysis. These techniques have been shown to perform fairly well in the context of whole-brain functional connectivity, particularly for acquisition artifacts (Ciric et al., 2017; Kassinopoulos and Mitsis, 2020a; Parkes et al., 2018; Xifra-Porxas et al., 2020). However, their performance on task-based studies or studies with limited field of view (e.g. Pattinson et al., 2009) has not been well addressed. Often the global signal (GS), defined as the mean timeseries across all voxels in the brain (or gray matter (GM)), is regressed out from the data as it is largely influenced by SLFOs (Birn et al., 2006; Chang and Glover, 2009a, 2009b; Falahpour et al., 2013; Golestani et al., 2015; Golestani and Chen, 2020; Kassinopoulos and Mitsis, 2019; Shmueli et al., 2007). However, this practice is somewhat controversial as there is evidence that GS variations may also reflect neuronal activity (Liu et al., 2017; Murphy and Fox, 2017; Power et al., 2017). Due to this, several methods that assess the effect of global signal regression (GSR) on connectivity measures have been proposed (Carbonell et al., 2014; Falahpour et al., 2018; Nalci et al., 2019b, 2019a), as well as alternative preprocessing approaches that avoid some of the limitations of GSR (Aquino et al., 2020; Carbonell et al., 2011; Glasser et al., 2018).

As the effectiveness and reliability of data-driven techniques is not yet fully understood, fMRI studies often employ model-based techniques that utilize concurrent external recordings (e.g. levels of CO_2_, continuous arterial blood pressure etc.) and, thus, are more directly related to specific sources of confounds without the risk of removing signal of interest. The physiological recordings most commonly acquired are the photoplethysmograph (PPG) and the respiratory bellow signal due to the availability of the necessary equipment in the majority of MR units and due to that the transducers are well tolerable for participants. PPG and the respiratory bellow capture cardiac and breathing activity, respectively, and are commonly used to account for high-frequency fMRI artifacts due to cardiac pulsatility (∼1 Hz) and breathing motion (∼0.3 Hz), using a technique termed RETROICOR. In addition, these two signals are often used with convolution models to account for SLFOs driven by variations in HR (Chang et al., 2009) and breathing patterns (Birn et al., 2008).

RETROICOR was proposed by Glover et al. (2000) and has been implemented in built-in toolboxes and external libraries of widely used fMRI analysis software packages such as FSL (Jenkinson et al., 2012), SPM (Kasper et al., 2017) and AFNI (Cox J.S., 1996). The component of RETROICOR related to cardiac pulsatility artifacts, referred to here as Card-RETROICOR, assumes that the pulse-related oscillations are phase-locked to the cardiac cycles. The cardiac cycles are considered to start and end at time intervals indicated by adjacent peaks in the PPG recording or adjacent R-waves in the electrocardiograph (ECG). Card-RETROICOR uses a Fourier series to define a cardiac pulsatility waveform (CPW) for each voxel timeseries, which is present in each cardiac cycle. Depending on how long the period of a cardiac cycle is, the CPW is extended or shrunk in time to achieve proper alignment with the cycle. From a systems theory perspective, Card-RETROICOR assumes that the pulse-related oscillations are described by a non-causal system as it requires knowledge of future input values (i.e. the timing of the following peak in PPG) to estimate the output at a specific timepoint. However, physiological systems are considered to be causal where the output depends on past and current inputs but not future inputs, and this raises the question whether a causal system can be formulated that can better model pulse-related oscillations.

In this study, we propose a refinement to the approach followed in Card-RETROICOR for modeling cardiac pulsatility that makes use of causal convolution models, inspired by their success in modeling BOLD fluctuations induced by variations in HR and breathing patterns (Birn et al., 2008; Chang et al., 2009), as well as changes in neuronal activity (Boynton et al., 1996). Specifically, we propose the cardiac pulsatility model (CPM) that describes pulse-related fluctuations in fMRI as the convolution of a CPW with a train of pulses located at the time of cardiac contractions. Essentially, Card-RETROICOR and CPM differ only in that the former assumes a CPW that is phase-locked to the cardiac cycle whereas the latter assumes a CPW of constant duration. Therefore, in the case of CPM, when two adjacent cardiac contractions occur close to each other compared to the typical time interval, the end of the first CPW overlaps with the beginning of the following CPW. As a result, the BOLD response, during this overlapping period, is effectively the sum of the individual waveforms, similar to the wave interference observed in nature with other types of waves. By setting the duration of the CPW equal to the average time interval between cardiac contractions, in the case of a recording with a constant HR, the CPWs of adjacent cardiac cycles do not overlap and can be in principle identical to the CPWs estimated with Card-RETROICOR. However, in the case that the HR exhibits pronounced fluctuations, even if the two models consider the same shape of CPW, their predicted outputs can have substantial differences. Using a cross-validation framework, we compare Card-RETROICOR with CPM in terms of variance explained in the fMRI BOLD signal. We hypothesized that CPM would outperform Card-RETROICOR, particularly for subjects with high HR variability (HRV).

Särkkä et al. (2012) have shown that a stochastic state space model, termed DRIFTER, outperforms RETROICOR in terms of variance explained, which was attributed to the fact that DRIFTER, in contrast to RETROICOR, is able to adapt to changes in shape and amplitude of cardiac and respiratory artifacts throughout time. While this finding suggests the presence of variations in the amplitude of pulse-related artifacts in fMRI, the underlying mechanisms of these variations cannot be determined with DRIFTER due to the data-driven nature of this model. To address this, in the present work, using a variant of CPM we also examine whether variations in the amplitude of fMRI pulse-related fluctuations can be attributed to variations in the amplitude of PPG pulses (PPG-Amp), that are believed to arise from systemic hemodynamic changes (e.g. changes in central venous pressure; Reisner et al. (2008)).

Furthermore, we examine the association between PPG-Amp and the SLFOs present in the GS. Even though recent sleep studies have demonstrated a strong association of PPG-Amp and GS (Özbay et al., 2019, 2018), it is still unclear whether the same relationship is present during resting conditions, and importantly whether fluctuations in PPG-Amp can assist in physiological noise correction. Previously, we have presented a framework for estimating scan-specific physiological response functions (PRFs) that are used to model the effect of HR and breathing pattern in SLFOs present in the GS (Kassinopoulos and Mitsis, 2019). By doing so, the SLFOs timeseries can be used as a nuisance regressor in the preprocessing of the fMRI data. Here, we investigate whether there is any benefit in considering PPG-Amp, in addition to HR and breathing activity, for modeling the effect of SLFOs on the GS.

Our results suggest that CPM explains more variance than Card-RETROICOR, particularly for subjects with high HRV. They also suggest that fluctuations in PPG-Amp do not covary with the amplitude of the cardiac-related pulses observed in fMRI but they can explain variance in the GS that is not captured by fluctuations of cardiac or breathing activity. Scripts for the models examined in the present study are available on git repository https://github.com/mkassinopoulos/Noise_modeling_based_on_PPG.

## 2. Methodology

Unless stated otherwise, the preprocessing and analysis described below were performed in Matlab (R2018b; Mathworks, Natick MA).

### 2.1 Human Connectome Project (HCP) Dataset

We used resting-state fMRI scans from the HCP S1200 release (Glasser et al., 2016; Van Essen et al., 2013). The HCP dataset includes, among others, T1-weighted (T1w) images and resting-state fMRI data (eyes-open and fixation on a cross-hair) from healthy young individuals (age range: 22-35 years) acquired on two different days. On each day, two 15-minute scans were collected, one with a left-right phase encoding (PE) direction and one with a right-left PE direction. fMRI acquisition was performed with a multiband factor of 8, spatial resolution of 2 mm isotropic voxels, and a repetition time (TR) of 0.72 s (Glasser et al., 2013).

The minimal preprocessing pipeline for the resting-state HCP dataset is described in Glasser et al. (2013). In brief, the pipeline included gradient-nonlinearity-induced distortion correction, motion correction, EPI image distortion correction and non-linear registration to MNI space. The motion parameters are included in the database for further correction of motion artifacts.

In the present work, we used the minimally-preprocessed data along with T1w images provided in volumetric MNI152 space. We considered 100 subjects who had good quality PPG and breathing activity (respiratory bellow) recordings in all four scans, as assessed by visual inspection.

### 2.2 Preprocessing and analysis of physiological signals

The detection of peaks in PPG with good time accuracy was important for the techniques examined in this study. The timings of the peaks in PPG were used to model high-frequency (∼1 Hz) cardiac pulsatility artifacts in fMRI and low-frequency (∼0.1 Hz) physiological fluctuations due to variations in HR reflected on the GS, while the amplitudes of the PPG peaks were used to model low-frequency fluctuations that may be unrelated to HR variations. Therefore, to facilitate peak detection, the PPG signal was initially band-pass filtered with a 2^nd^ order Butterworth filter between 0.3 and 10 Hz. The minimum peak distance specified for peak detection varied between 0.5 and 0.9 s, depending on the subject’s average HR. For a given set of detected peaks, the HR signal was computed in beats-per-minute (bpm) by multiplying the inverse of the time differences between pairs of adjacent peaks with 60, and evenly resampled at 10 Hz. The amplitudes of the peaks, referred to later as photoplethysmographic amplitude (PPG-Amp), were also evenly resampled at 10 Hz. The resampling of HR and PPG-Amp was done using linear interpolation. Note that several subjects in HCP demonstrated a constant HR of 48 bpm, which is likely due to erroneous PPG recording. None of these subjects were considered in our study. Furthermore, several scans considered here illustrated HR traces with outliers at different timepoints. The values of HR at the timepoints of outliers were corrected using linear interpolation (for more information see Methods in (Kassinopoulos and Mitsis, 2019)).

The breathing signal was detrended linearly and corrected for outliers using a median filter in a similar manner to Kassinopoulos and Mitsis (2019). Subsequently, the breathing signal was low-pass filtered at 5 Hz with a 2^nd^ order Butterworth filter and z-scored.

### 2.3 Preprocessing of fMRI data

fMRI data as well as 16 nuisance regressors were first high-pass filtered at 0.008 Hz. The nuisance regressors consisted of the 6 motion parameters along with their derivatives and 4 regressors related to breathing motion (i.e. 2^nd^ order RETROICOR using only the regressors related to breathing motion). Subsequently, the nuisance regressors were removed from the fMRI data. To extract the GS of each scan, we initially performed tissue segmentation on the T1w images in the MNI152 space using FLIRT in FSL 5.0.9, which generated probabilistic maps for the gray matter (GM), white matter (WM) and cerebrospinal fluid (CSF) compartments (Zhang et al., 2001). Afterwards, the GS was calculated by estimating the mean timeseries across all voxels with a probability of belonging to GM above 0.25. The choice of the threshold value was made based on visual inspection while overlaying the probabilistic maps of GM on the T1w images.

### 2.4 High-frequency cardiac fluctuations

#### 2.4.1 Cardiac-related RETROICOR (Card-RETROICOR)

RETROICOR stands for RETROspective Image-based CORrection and is a technique proposed by Glover et al. (2000) for removing high-frequency fluctuations due to cardiac pulsatility and breathing motion. While similar steps are performed to remove the aforementioned confounds, different mechanisms underly them. In this study, we compared the variance explained in fMRI data using Card-RETROICOR with the variance explained using CPM which is a physiologically plausible model for pulsatility artifacts.

To obtain the Card-RETROICOR regressors we followed the following three steps:

1. The cardiac phase was first defined based on the timings of the PPG peaks using the relation:

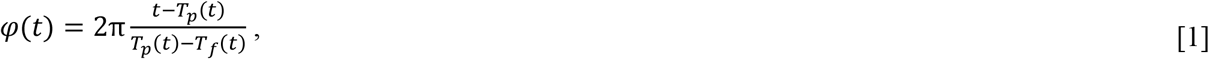

where *T*_*p*_(*t*)and *T*_*f*_(*t*)indicate the time of the nearest peaks from past and future timepoints, respectively.

2. A basis set of cosines and sines was created as follows:

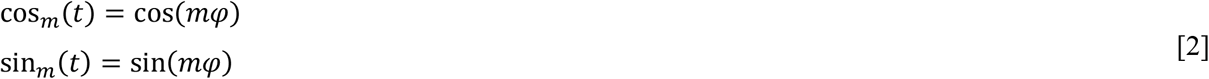

where *m* is the order of the Fourier basis set. Typically, a 2^nd^ order Card-RETROICOR is employed, which corresponds to four nuisance regressors. It has been suggested that whereas higher orders improve the fit, they carry the risk of overfitting the data (Harvey et al., 2008). However, as the optimal order may depend on parameters of the fMRI pulse sequence, such as the duration of scan and TR, one of the goals of this study was to determine the optimal order for Card-RETROICOR and CPM when applied to the resting-state fMRI data of HCP (see Section 2.4.4).

3. Finally, the nuisance regressors cos_*m*_(*t*)and sin_*m*_(*t*)were downsampled to match the fMRI acquisition rate, yielding the regressors cos_*m*_[*n*] and sin_*m*_[*n*]. Note that we use parentheses and brackets to distinguish the original and low-sampled timeseries, and not for distinguishing continuous and discrete signals.

The resulting nuisance regressors were used in the design matrix of the general linear model (GLM) in order to model the pulse-related artifacts in each voxel timeseries. Through linear regression in GLM, each nuisance regressor is assigned with a beta parameter *β*. In the case that only cardiac-related regressors are included in the design matrix, the voxel timeseries can be represented as:

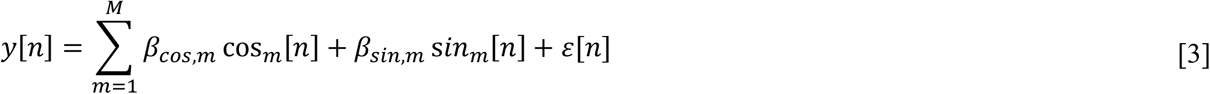

where *ε*[*n*] corresponds to random Gaussian errors. Section 2.4.4 describes how the fit of Card-RETROICOR regressors on the data was assessed and compared with CPM. Note that the beta parameters essentially define the cardiac pulsatility waveform (CPW) during a cardiac cycle. Specifically, CPW is obtained using the relation:

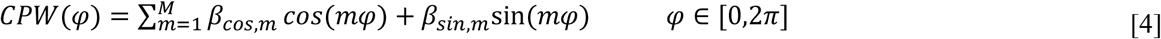

#### 2.4.2 Cardiac pulsatility model (CPM)

Here, we propose an alternative model for capturing pulse-related artifacts in fMRI, termed cardiac pulsatility model (CPM). CPM assumes that pulse-related artifacts can be modeled as the output of a causal linear time-invariant (LTI) system, where the input is a train of pulses corresponding to cardiac contractions. Therefore, to describe these artifacts we employed the convolution representation:

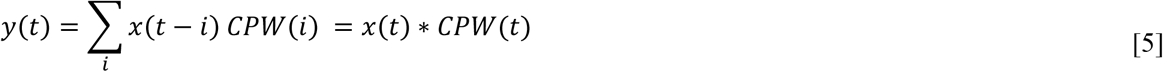

where *x*(*t*)is the input of the model reflecting the cardiac contractions and *CPW*(*t*)is the cardiac pulsatility waveform and has the role of the model impulse response. The timepoints of the cardiac contractions were considered to coincide with the timings of the PPG peaks. Two different inputs were considered that led to the examination of two variants of CPM, namely the CPM_CA_ and CPM_VA_ model: the first input, denoted by *x*_*CA*_(*t*), is based on the assumption that all pulses had a constant amplitude of one, while in the second, denoted by *x*_*VA*_(*t*), the amplitudes of the pulses were equal to the amplitudes of the PPG peaks. Both inputs were defined at the PPG’s sampling rate (400 Hz). The pulses consisted of one sample duration each with a constant (unit value) or varying amplitude, depending on the input (i.e. *x*_*CA*_(*t*)or *x*_*VA*_(*t*)), and all remaining samples were equal to zero. The impulse response *CPW*(*t*)was defined as a linear combination of modified Fourier basis functions of one cycle length as shown below:

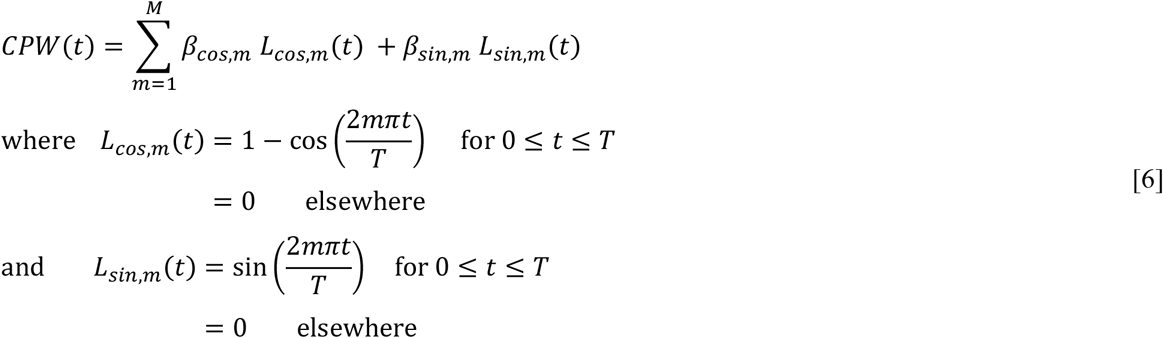

The fundamental period *T* varied on a scan basis and was equal to the average cardiac cycle duration within a scan (i.e. average time difference across pairs of adjacent PPG peaks). The cosine terms were subtracted from one so that all basis functions begin and end at zero. This mathematical manipulation ensures the physiological plausibility of the model as it leads to impulse responses that have zero amplitude at time zero and at times larger that *T*. Note that based on the properties of convolution, Eq. 5 can also be expressed as:

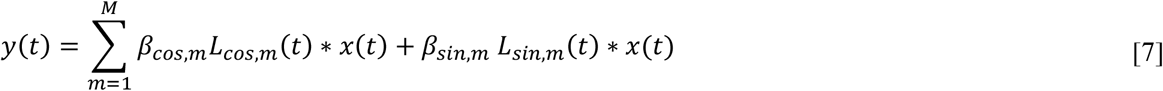

As in Card-RETROICOR, nuisance regressors extracted from the CPM were subsequently downsampled to the fMRI acquisition timeline and used in the GLM design matrix. Suppl. Fig. 1 illustrates the main steps for obtaining the pulse-related nuisance regressors and the CPWs using CPM and the two variants of CPM (for a demonstration of how these steps can be implemented in Matlab, see script RETR_CPM_one_voxel.m on repository https://github.com/mkassinopoulos/Noise_modeling_based_on_PPG).

**Fig. 1.**
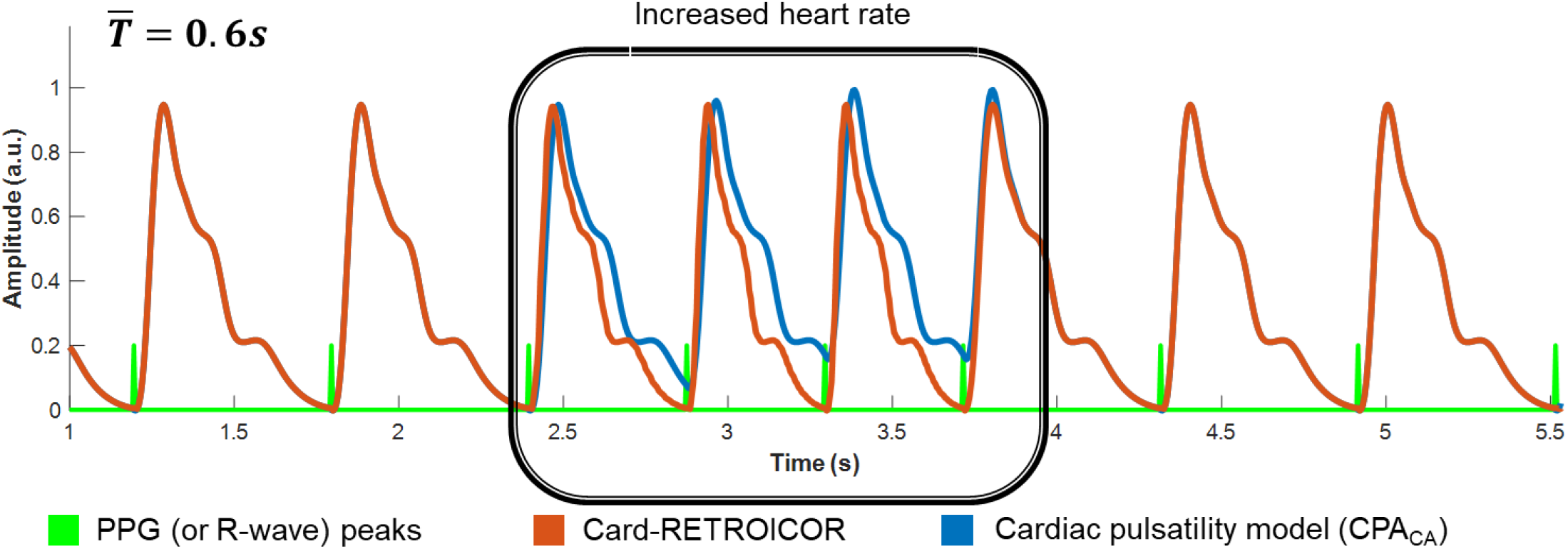
Schematic illustration of the output of Card-RETROICOR and Cardiac pulsatility model (CPA_CA_) during a period with increased heart rate (HR). Card-RETROICOR and CPA_CA_ use a linear combination of Fourier basis functions to construct the cardiac pulsatility waveform (CPW) and model the pulse-related BOLD fluctuations. When the same model order is used and the HR (time difference between adjacent PPG peaks) during a scan is fairly constant, the two models yield similar CPWs. In addition, the output of the two models is also similar during periods with HR close to the average HR. However, as illustrated at the center of the plot, RETROICOR generates the output based on the cardiac phase, therefore, during segments with increased HR, the CPW is condensed in time to achieve proper alignment with the cardiac cycle (similarly, although not shown here, during periods with decreased HR, the CPW in RETROICOR is extended). In contrast, CPM_CA_ assumes that the duration of the CPW is constant and during period with increased HR, the CPWs of two adjacent cardiac cycles may overlap with each other.

Even though both RETROICOR and CPM employ the Fourier basis functions to construct a CPW for each voxel, RETROICOR assumes that the pulse-related artifacts are phase-locked to the cardiac cycles whereas CPM assumes that these artifacts are time-locked to the cardiac contractions. As a consequence, the two models exhibit slightly different outputs at periods with increased or decreased HR. The simulated model outputs in Fig. 1 illustrate this difference for the case of a transient increase in HR.

Various MRI techniques have been proposed to measure intracranial CPWs (e.g. phase-contrast, magnetic resonance encephalography, gradient-recalled-echo MRI) as these waveforms are considered a useful biomarker in certain cerebrovascular diseases (Bianciardi et al., 2016; Rajna et al., 2021; Wagshul et al., 2011; Whittaker et al., 2021), and are often studied in order to understand the role of intracranial cardiac pulsatility in the glymphatic activity (Fultz et al., 2019; Kiviniemi et al., 2016). With this in mind, here we sought to investigate whether the CPM applied to BOLD fMRI data may provide an alternative technique for measuring intracranial pulsatility. Specifically, we sought to investigate the consistency of CPWs across subjects.

To extract CPWs averaged across subjects, we followed the following steps for the two variants of CPM (i.e. CPM_CA,_ CPM_VA_): For a given scan, we estimated the beta parameters of Eq. 7 through linear regression considering a period *T* equal to the average cardiac cycle duration of the scan. Subsequently, we reconstructed the CPW within each voxel based on Eq. 6, using the estimated beta parameters and a period *T* of 1 s, as well as a time-step of 0.025 s. Note that even though a varying period *T* was considered for the estimation of the beta parameters, a normalized constant period was used for the reconstruction of the group average CPW, since the waveform is of interest here rather than the exact duration. Finally, the amplitude of the CPW in each voxel was normalized to a maximum absolute value of one and multiplied by the correlation coefficient related to the variance explained by CPM in the associated voxel. In this way, CPWs in regions that were more prone to pulse-related fluctuations had larger intensity amplitudes. Finally, the CPWs were averaged across subjects considering only scans from the first session and the same PE direction.

#### 2.4.3 Time alignment of pulse-related regressors and fMRI data

When the heart contracts, a pulse pressure wave propagates from the heart through the vasculature to the whole body giving rise to cardiac pulses in the arteries that are captured, among other hemodynamic signals, in the PPG. The vascular path between heart and finger, where PPG is typically recorded, may differ in terms of travelling distance compared to the path between heart and arteries in the brain vasculature. As a result, a pulse originating from the heart will typically reach the finger at a different time compared to an artery in the brain. This time difference should be ideally incorporated in the analysis when extracting pulse-related regressors. To examine whether a time difference should be used in the HCP fMRI data that can improve the performance of the examined models, we repeated the analysis with each of the models considered in this study for lag times varying from −2 s to 2 s in steps of 0.1 s. In practice, to account for a specific lag time, the PPG signal depending on the lag time examined was shifted towards negative or positive times before extracting the pulse-related regressors. Section 2.4.4 describes how CARD-RETROICOR and the two variants of CPM (i.e. using inputs *x*_*CA*_(*t*)and *x*_*VA*_(*t*)) were compared as well as how the optimal lag time was determined for each of the models.

#### 2.4.4 Comparison of models for pulse-related fluctuations

To compare the performance between Card-RETROICOR, CPM_CA_ and CPM_VA_, a 3-fold cross-validation framework was employed to avoid overfitting as the models differed in flexibility (number of free parameters). Each fMRI scan was partitioned into three segments of about 5 min each. One segment was used as the validation dataset for assessing the performance of the model and the remaining two segments were used as the training dataset. This step was repeated three times with each of the three segments used exactly once as the validation data. In each fold, the voxel-specific beta parameters in Eq. 3 (Card-RETROICOR) and Eq. 7 (CPM) were estimated from the training dataset and, subsequently, used in the validation dataset to model the pulse-related fluctuations. The goodness-of-fit was assessed based on the correlation between the output of the model and the voxel timeseries in the validation dataset. Subsequently, the mean correlation across the three folds was calculated. To compare two models, the means correlations averaged across all voxels in the brain and all four scans of each subject were first computed and, then, a paired *t*-test was performed based on the mean correlation values of the 100 subjects for the two models.

Apart from the performance of each model type in terms of goodness-of-fit, we also examined the optimal model order of Fourier series and the optimal lag time as these two parameters may have significant impact on the performance. Note that before comparing model types, the model order and lag time had to be first optimized as these two parameters have a significant impact on the performance. To this end, the model performance was initially assessed for each of the three model types for model order varying from 1 to 8 and lag time varying from −2 s to 2 s in steps of 0.1 s. The comparison of the three models was subsequently made using for each model the order and lag time that yielded the best performance across all subjects.

We also repeated the comparison between the models considering the PPG raw signal as the output target rather than the voxel timeseries. This comparison allowed us to illustrate how well the three models capture cardiac pulsatility, including the low-frequency variations in the envelope of the pulses, on a timeseries that, in contrast to the fMRI data examined here, has a high sampling-rate (400 Hz) and thus does not suffer from aliasing. Note that the comparison of the models with regards to their fit on the PPG was made to give us insight into what features of the output timeseries these models are able to capture. However, the choice for the more appropriate model in the context of fMRI was done based on the model fit in the voxel timeseries.

To assess regional variability in the performance of the pulse-related models, statistical maps were generated. For visualization purposes, maps shown here were overlaid on structural images after being registered to the MNI152 space (1 mm spatial resolution) using FSL’s FLIRT registration tool (Jenkinson and Smith, 2001), as incorporated in the MANGO software (Lancaster, Martinez; www.ric.uthscsa.edu/mango).

### 2.5 Systemic low-frequency oscillations (SLFOs)

Apart from removing high-frequency cardiac and breathing fluctuations, physiological recordings can also be used to yield nuisance regressors that account for the effect of SLFOs (Tong et al., 2019). A common approach for removing SLFOs is to use convolution models where the inputs are the HR and a breathing-related variable (e.g. respiration volume per time; RVT) and the impulse responses are the so-called cardiac (CRF) and respiration (RRF) response functions (Birn et al., 2008; Chang et al., 2009). The outputs of these convolution models are subsequently used as nuisance regressors in the GLM. The physiological response functions (PRFs), RRF and CRF, have been shown to vary across subjects (Falahpour et al., 2013) and, to a less degree, across sessions within-individuals (Kassinopoulos and Mitsis, 2019), which led to the use of the GS of a scan as the model output for estimating subject- and scan-specific PRFs.

Apart from fluctuations in HR and breathing patterns, the GS has been recently shown to be linked, during sleep, to fluctuations in PPG-Amp (Özbay et al., 2019, 2018). However, as PPG-Amp is affected by various physiological processes such as breathing, cardiac activity and arterial blood pressure (Reisner et al., 2008), it is unclear whether fluctuations of PPG-Amp explain an additional fraction of GS variance compared to HR and breathing pattern variations.

In this study, we used cross-correlation analysis to better understand the relation between the physiological sources (HR, breathing patterns and PPG-Amp) and the GS during resting conditions. Fluctuations in breathing patterns were quantified using the respiration volume (RV) measure, which is defined as the standard deviation of the breathing signal within a sliding window of 6 s (Chang et al., 2009). The GS was upsampled to 10 Hz to match the sampling rate of physiological variables before proceeding with the cross-correlation analysis.

In addition, we employed the framework proposed in Kassinopoulos and Mitsis (2019) to examine whether a convolution model associated to PPG-Amp variations may explain variance in the GS not attributed to HR and breathing variations. Specifically, we examined whether considering a PRF associated to PPG-Amp variations, (referred to below as PPG-Amp response function; PARF) along with the components related to HR and breathing pattern variations can improve the variance explained on the GS compared to the standard approach of considering only HR and breathing patterns. The nuisance regressors related to HR, PPG-Amp and RV were defined as follows:

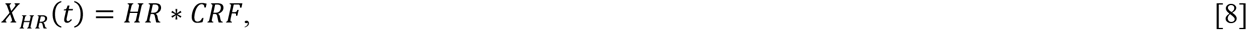

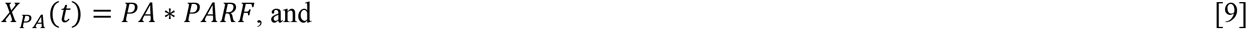

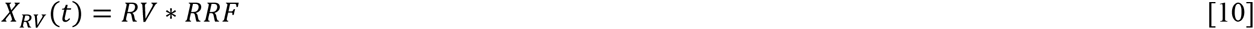

where *PA* is the PPG-Amp. We considered the double gamma function (i.e., the sum of two gamma functions) to construct the *PRF* curves. The gamma function is defined as:

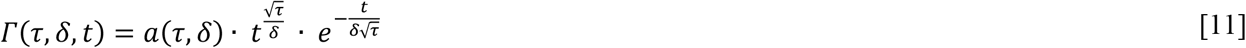

where the parameters *τ* and *δ* indicate the (approximate) time of peak and dispersion of the function, and the parameter *α* is a scaling factor which normalizes the peak value of the gamma function to 1. The *PRF* curves were defined as follows:

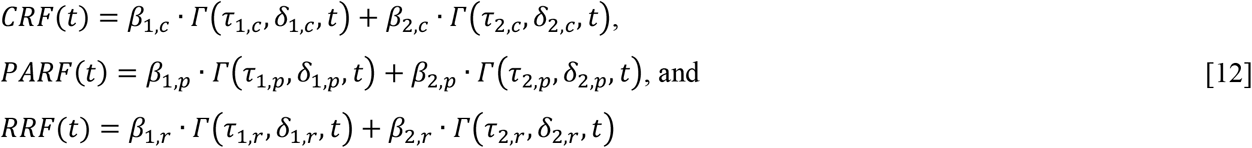

The procedure for estimating scan-specific PRFs is described in detail in Kassinopoulos and Mitsis (2019). In brief, the parameters of the PRFs for a given scan were first estimated using a genetic algorithm (GA) implemented in Matlab R2018b’s Global Optimization Toolbox. The parameter vectors ***τ*** (*τ*_1,*c*_, *τ*_2,*c*_, *τ*_1,*p*_, *τ*_2,*p*_, *τ*_1,*r*_, *τ*_2,*r*_) and ***δ*** (*δ*_1,*c*_, *δ*_2,*c*_, *δ*_1,*p*_, *δ*_2,*p*_, *δ*_1,*r*_, *δ*_2,*r*_) were bounded between 0-20 seconds and 0-3 seconds, respectively. A stopping criterion of 100 generations was set, as it was found to be adequate for convergence. GA searches in the parameter space defined by the boundaries to estimate the parameters that maximize the objective function (Pearson correlation coefficient). Specifically: 1. for a set of given parameters, the three *PRF* curves were constructed. Subsequently, 2. the HR, PPG-Amp and RV signals (sampled at 10 Hz) were convolved with *CRF, PARF* and *RRF*, respectively, to extract the corresponding nuisance regressors and then downsampled to match the fMRI acquisition rate. 3. The beta parameters were estimated through linear regression (GLM), whereby the GS was the dependent variable and the three nuisance regressors were the three explanatory variables. 4. Finally, the Pearson correlation coefficient between the GS and the model prediction was calculated. The estimated parameter values yielded by the GA were subsequently refined using the interior-point gradient-based algorithm with a stopping criterion of 100 maximum iterations (implemented in Matlab R2018 as well).

To obtain the main trend of the PRFs observed in the examined population, the estimated PRFs were averaged across all scans (*N*_*scans*_ = 400) using weighted averaging as shown below:

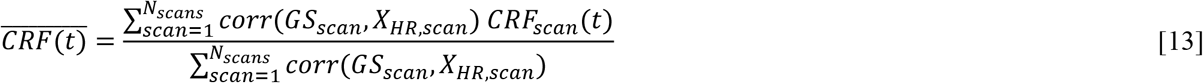

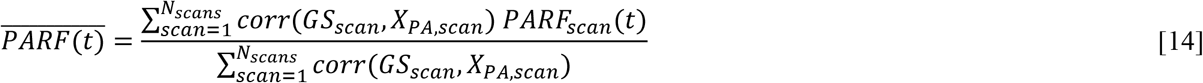

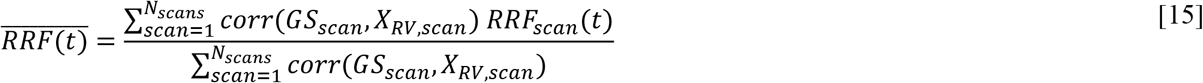

where the correlation values between *GS*_*scan*_ and the associated nuisance regressors (*X*_*HR,scan*_, *X*_*PA,scan*_ and *X*_*RV,scan*_) found for each scan were used as the weighting coefficient.

To assess the potential benefit of considering a regressor associated to PPG-Amp variations in physiological noise modeling, we compared two models for extracting the SLFOs using the GS. The standard model quantifies the effect of SLFOs in the GS using only the variations in HR and breathing patterns, whereas the proposed model uses HR, breathing patterns and PPG-Amp. The two models can be expressed as follows:

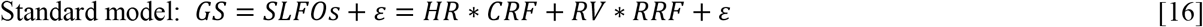

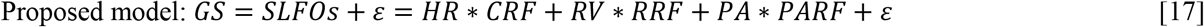

where *ε* corresponds to random Gaussian errors. Similar to the pulse-related models, we employed a 3-fold cross-validation framework to compare the performance of the standard and proposed model. The GS of each scan was partitioned into three segments of about 5 min each. One segment was used as the validation dataset for assessing the performance of the model and the remaining two segments were used as the training dataset. This step was repeated three times with each of the three segments used exactly once as the validation dataset. The first or last 42 timepoints (∼30 seconds) of the segments considered in the training dataset were excluded if they were adjacent to the boundaries of the validation dataset. For instance, in the cases that the validation dataset consisted of the second segment and the first and third segment consisted the training dataset, the last 42 timepoints of the first segment and the first 42 timepoints of the third segment were excluded from the subsequent analysis. The exclusion of the boundaries was done to prevent data leakage between the training and validation dataset that may arise due to high autocorrelation in the fMRI signal, as this may bias the estimation of the PRFs and inflate model performance. This step was not needed for the pulse-related models as the memory of these models is at around one second and therefore the data leakage at the boundaries of the 5 min segments can be considered negligible. In each fold, scan-specific PRFs were estimated from the training dataset and, subsequently, used in the validation dataset to model the SLFOs. The goodness-of-fit was assessed based on the correlation between the GS and estimated SLFOs in the validation dataset. Finally, the mean correlation across the three folds was calculated. To compare the standard with the proposed model, the mean correlations of the four scans of each subject were first averaged and, subsequently, a paired *t*-test was performed based on the mean correlation values of the 100 subjects for the two models.

## 3. Results

The present study examined three models for capturing fMRI fluctuations related to cardiac pulsatility, namely Card-RETROICOR, CPM_CA_ and CPM_VA_, as well as two models for obtaining the SLFOs from fluctuations in the GS. While the repetition time in the HCP fMRI data (TR=0.72 s) is relatively short compared to typical fMRI data, it is still not short enough to avoid aliasing of pulse-related fluctuations (Suppl. Fig. 1). In addition, the aliased frequency location varies considerably across individuals due to differences in HR making the visual inspection of the model fit in fMRI data more difficult (Suppl. Fig. 2). Therefore, to better understand the capabilities of the three pulse-related models, the models were first compared with regards to their ability in modeling the high-frequency oscillations of the PPG signal, which, due to its relatively high sampling rate (400 Hz), did not exhibit any aliasing. The comparison of the three pulse-related models in modeling high-frequency PPG oscillations is presented in Section 3.1, whereas Section 3.2 presents the comparison of the models with regards to the variance explained in fMRI voxel timeseries. Section 3.2 also examines the relation of pulse-related fMRI fluctuations with the direction of phase encoding in fMRI acquisition. Finally, Section 3.3 examines the relation of the low-frequency oscillations in PPG-Amp with the SLFOs observed in GS, which are known to be driven, among others, by variations in HR and breathing patterns (Birn et al., 2006; Kassinopoulos and Mitsis, 2019; Shmueli et al., 2007).

**Fig. 2.**
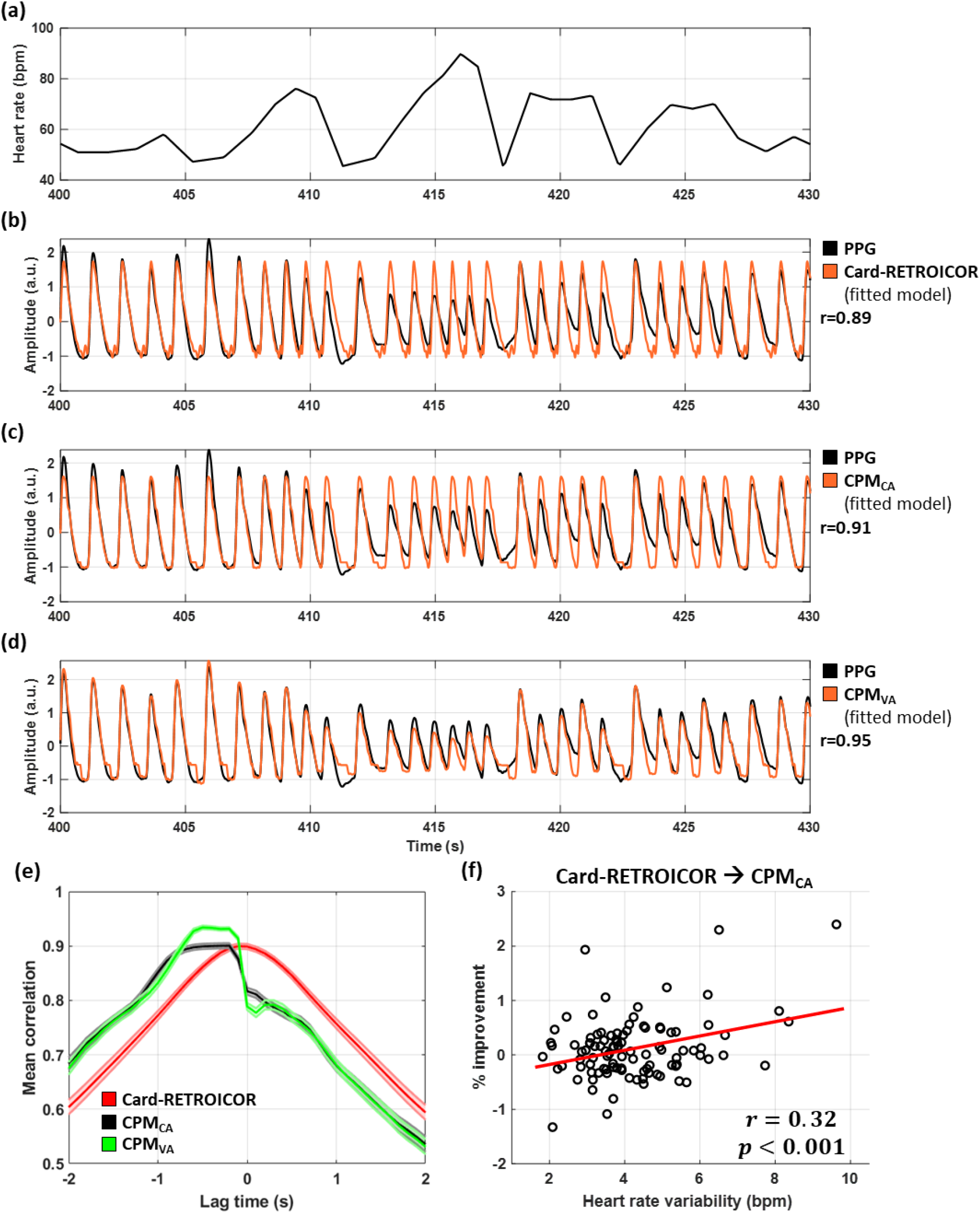
Model fit of pulse-related models for PPG. **(a)** Trace of HR of a subject with high heart rate variability (HRV; S159138-R1LR) during a 30 s time segment with pronounced fluctuations. **(b)-(d)** PPG model fit for Card-RETROICOR, CPM_CA_ and CPM_VA_. All three models captured the high-frequency (∼1 Hz) fluctuations related to cardiac pulsatility. However, only CPM_VA_, which incorporates the low-frequency (∼0.1 Hz) fluctuations in PPG-Amp was able to represent these fluctuations in the PPG. **(e)** Cross-correlation averaged across subjects for Card-RETROICOR, CPM_CA_ and CPM_VA_. CPM_VA_ exhibited significantly better performance compared to the other two models as it accounted for the low-frequency fluctuations of PPG. Note that while Card-RETROICOR and CPM_CA_ exhibited similar maximum mean cross-correlation values, the peak of the latter was broader (∼0.5 s) compared to the peak of the former (∼0.1 s). **(f)** Relative percentage (%) improvement achieved by the proposed model CPM_CA_ compared to Card-RETROICOR. The relative percentage improvement for a given scan was defined as 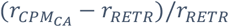 where *r*_*RETR*_ and 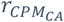 are the correlation values obtained for models Card-RETROICOR and CPM_CA_, respectively. Even though the model that yielded the best goodness-of-fit varied across scans, scans that were characterized by larger HRV (*p*<0.001) tented to show better goodness-of-fit with CPM_CA_ than with Card-RETROICOR. The comparison was performed between these two models as these models exhibited better performance than CPM_VA_ when considering the variance explained in the fMRI data (Fig. 4).

### 3.1 High-frequency oscillations in the photoplethysmographic (PPG) signal

All three models examined for pulse-related oscillations were able to model the effect of cardiac pulsatility in the PPG. When considering a 3-fold cross-validation framework, the three models showed improved performance for higher order Fourier series, reaching a plateau at about 4^th^ order. Considering the highest examined (8^th^) order, which demonstrated the best performance, we observed a maximum cross-correlation averaged across subjects at time lags between −0.1 s and −0.5 s (Fig. 2e). Card-RETROICOR, CPM_CA_ and CPM_VA_ yielded maximum cross-correlation values of 0.900, 0.901 and 0.934, respectively. CPM_VA_ which, in contrast to RETROICOR and CPM_CA_, takes into account variations in PPG-Amp, yielded significantly better performance compared to the other two models (p<10^−31^), while Card-RETROICOR and CPM_CA_ yielded similar performance. Fig. 2a-d shows the goodness-of-fit for the three models when applied to the PPG of a subject that exhibited pronounced HR variations (the 3-fold cross validation framework was omitted for this figure). As we can see, all three models explained well the variations in the PPG signal overall, although only CPM_VA_ was able to capture the low-frequency fluctuations of the PPG-Amp. Apart from the absence of low-frequency fluctuations in the case of Card-RETROICOR and CPM_CA_, we were not able to visually observe any other differences in the output waveforms of the three models. However, as shown later, in the case of fMRI data, Card-RETROICOR and CPM_CA_ exhibited better performance compared to CPM_VA_. Due to this, we further tested whether scans with larger HRV values, where HRV was defined as the standard deviation of the HR signal, showed relative improvement in the case of CPM_CA_ as compared to Card-RETROICOR. For this test, an 8^th^ order model was used with the optimal lag time for each model. As can be seen in Fig. 2f, CPM_CA_ explained a larger fraction of variance for some scans compared to Card-RETROICOR, while the opposite was observed for other scans. Even though none of the two models was found to outperform the other one for all scans, we observed a trend for scans with high HRV to be characterized by higher relative improvement (Fig. 2f; r=0.32, p<0.001).

### 3.2 Effects of cardiac pulsatility in BOLD fMRI

Fig. 3a-c shows the correlation averaged across subjects (as well as across all voxels and scans in each subject) for the three models when the target output was the fMRI timeseries. As many voxels are not prone to pulse-related fluctuations, the correlation values averaged across all voxels were relatively low. For all model orders examined, Card-RETROICOR exhibited a peak in cross-correlation at lag time of −0.4 s, whereas the two CPM models exhibited a peak at lag time of −0.9 s (a negative lag time indicates that the PPG signal was shifted backward in time). To ease the comparison between models, Fig. 3d shows the mean correlation for all models along with the associated standard error corresponding to these optimal lag times. We observe that, in fMRI data, CPM_VA_ yielded poorer performance compared to Card-RETROICOR and CPM_CA_. Moreover, for all three models, the highest performance was achieved with a 6^th^ order Fourier series, whereas higher orders yielded slightly lower mean correlation values.

**Fig. 3.**
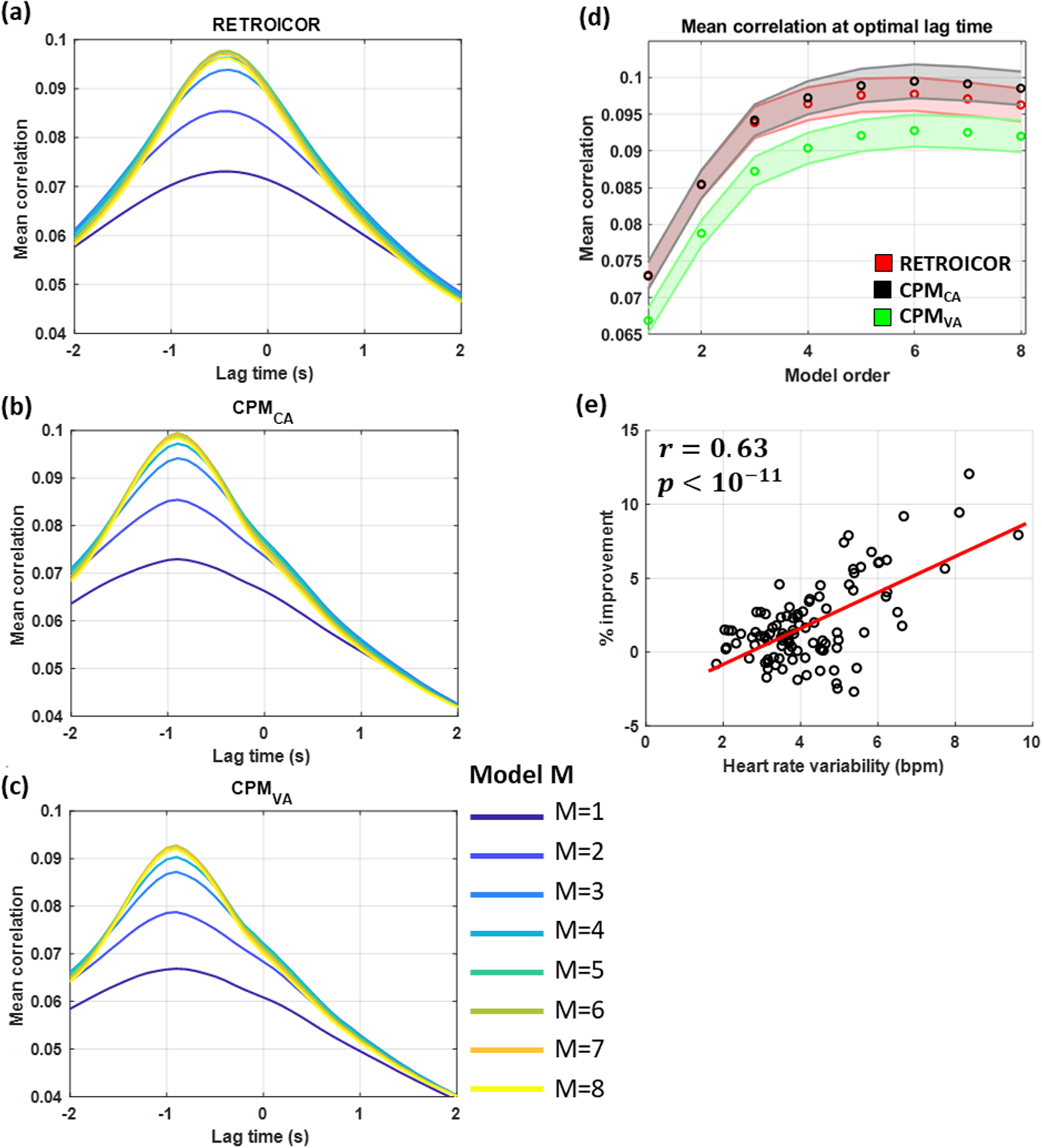
Performance of pulse-related models in fMRI data. (a)-(c) Correlation averaged across subjects for Card-RETROICOR, CPM_CA_ and CPM_VA_, respectively. Colors from blue to yellow indicate performance for higher model orders. For all three models, a unimodal curve was observed with highest mean correlations for lag times between −0.4 and −0.9 s. The optimal lag time for each model was consistent across model orders. Note that Card-RETROICOR exhibited slightly different optimal lag time compared to CPM_CA_ and CPM_VA_ which is likely due to differences in the cosine terms of the basis sets used in the three models. (d) Mean correlation with respect to model order for optimal lag time of −0.4 s for Card-RETROICOR and −0.9 s for the two variants of CPM. All models yielded a maximum mean correlation for a 6^th^ order model. In contrast to the model fit on PPG (Fig. 2e), incorporating the low-frequency fluctuations of PPG-Amp in the input signal was found to decrease the model performance (i.e. CPM_CA_ vs CPM_VA_). (e) Relative percentage (%) improvement with respect to HRV when comparing the proposed model CPM_CA_ with Card-RETROICOR. The relative percentage improvement for a given scan was defined as 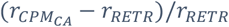 where *r*_*RETR*_ and 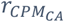 are the correlation values obtained for models Card-RETROICOR and CPM_CA_, respectively. The improvement achieved with CPM_CA_ compared to Card-RETROICOR was found to be correlated to the amplitude of HR fluctuations (p<10^−11^). Note, however, that for some of the scans with low HRV (<6 bpm), Card-RETROICOR yielded better goodness-of-fit than CPM_CA_.

Table 1 summarizes the performance of four specific models in order to illustrate the main steps that can lead to an improvement in the variance explained in the fMRI data. Model M1 corresponds to the 2^nd^ order Card-RETROICOR that is commonly employed in fMRI studies without considering any lag time, whereas model M2 considers the optimal lag time of Card-RETROICOR (−0.4 s). A paired *t*-test showed that considering the optimal lag time when extracting the nuisance regressors leads to a significant improvement in the variance explained (*p*<10^−24^). Moreover, employing a 6^th^ instead of a 2^nd^ order Fourier series resulted also in an additional improvement (*p*<10^−29^). Finally, when the optimal model order and lag time were considered, further improvement was achieved when modeling cardiac pulsatility using the proposed model CPM_CA_ compared to Card-RETROICOR (*p*<10^−8^). As hypothesized, when comparing CPM_CA_ (6^th^ order and −0.9 s) to Card-RETROICOR (6^th^ order and −0.4 s lag time), larger HRV values were associated with a larger relative improvement (Fig. 3e; r=0.63; *p*<10^−11^).

**Table 1.**
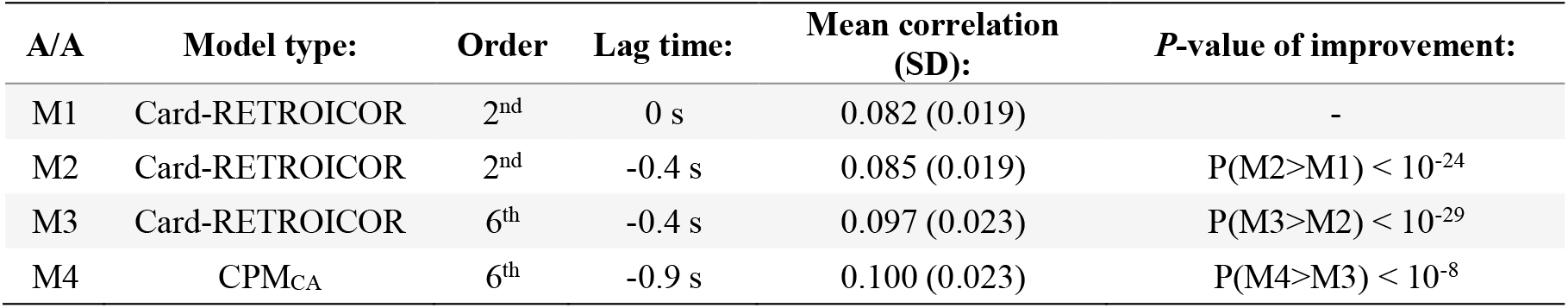
Comparison of pulse-related models in terms of variance explained in fMRI data

| A/A | Model type: | Order | Lag time: | Mean correlation (SD): | *P*-value of improvement: |
| --- | --- | --- | --- | --- | --- |
| M1 | Card-RETROICOR | 2<sup>nd</sup> | 0 s | 0.082 (0.019) | - |
| M2 | Card-RETROICOR | 2<sup>nd</sup> | -0.4 s | 0.085 (0.019) | $P(M2>M1) < 10^{-24}$ |
| M3 | Card-RETROICOR | 6<sup>th</sup> | -0.4 s | 0.097 (0.023) | $P(M3>M2) < 10^{-29}$ |
| M4 | CPM<sub>CA</sub> | 6<sup>th</sup> | -0.9 s | 0.100 (0.023) | $P(M4>M3) < 10^{-8}$ |

Fig. 4 shows correlation maps averaged across subjects (only scans with left-right PE direction from the first session were included) as obtained with Card-RETROICOR and CPM_CA_ when considering 6^th^ order Fourier series and the optimal lag time for each model (i.e. −0.4 s and −0.9 s for Card-RETROICOR and CPM_CA_, respectively). It also shows *t*-score maps derived with paired *t*-tests, indicating brain regions with significant differences in goodness-of-fit between the two models. As expected, both Card-RETROICOR and CPM_CA_ explained substantial variance in areas with CSF (e.g. areas around the brainstem, in the 4^th^ ventricle and the superior sagittal sinus) as well as in lateral sulcus and occipital cortex. At a significance level of *p*<0.01, CPM_CA_ demonstrated better performance in terms of variance explained, particularly in the occipital cortex, lateral sulcus and superior sagittal sinus, while none of the regions were associated with a better fit for Card-RETROICOR. When the significance level was set to higher *p* values, Card-RETROICOR was found to explain a larger fraction of variance in small clusters of voxels within the lateral ventricles.

**Fig. 4.**
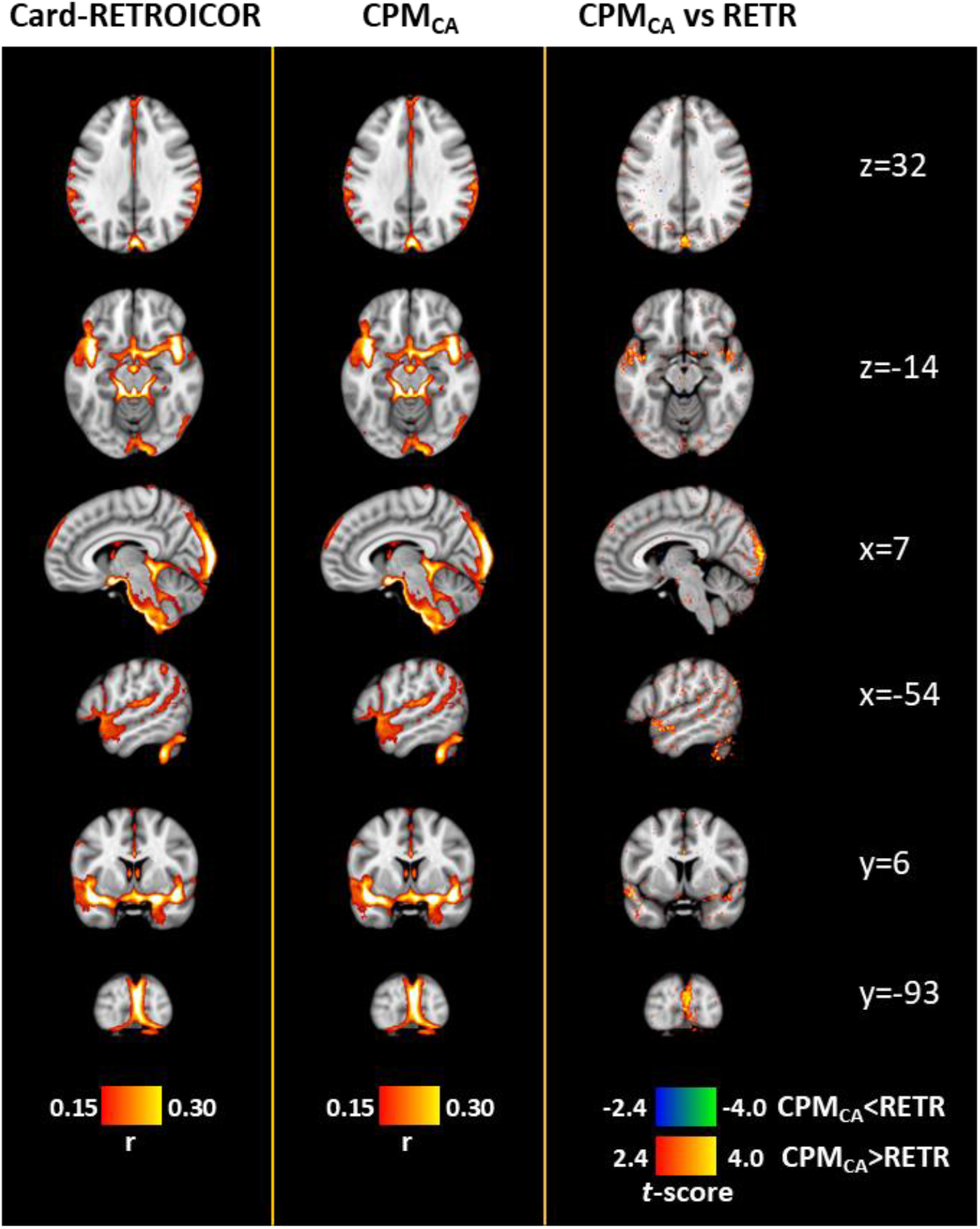
Correlation maps for pulse-related models averaged across all subjects (N=100; only scans with left-right PE directions from the first session were included here). The first and second columns show the variance explained using Card-RETROICOR and the proposed model CPM_CA_, respectively, while the third column shows *t*-stat maps indicating the areas with significant differences in correlation values between the two models (*p*<0.01). The correlation threshold value was chosen so that only the regions more susceptible to cardiac pulsatility are visible. Both Card-RETROICOR and CPM_CA_ revealed regions close to the basilar and vertebral arteries, in the 4^th^ ventricle, in the superior sagittal sinus, in the lateral sulcus and in the occipital lobe as being more prone to pulse-related oscillations. However, based on the *t*-stat map, CPM_CA_ explained a significantly larger fraction of variance than Card-RETROICOR in the occipital cortex, the lateral sulcus and superior sagittal sinus. With the level of significance set at *p*<0.01, no region was found to yield higher correlation values for Card-RETROICOR than for CPM_CA_, while for larger values of *p*-value, only the lateral ventricles show small clusters of regions where Card-RETROICOR outperformed CPM_CA_ in terms of variance explained. The statistical maps shown in this figure are available on: https://neurovault.org/collections/DHFETQTN/.

In a recent study, we investigated the role of physiological processes in fMRI connectome-based subject discriminability (Xifra-Porxas et al., 2020). One of our findings was that the connectome signature driven by cardiac pulsatility differs between left-right and right-left PE direction (see for example Fig. 5 in Xifra-porxas et al. (2020)). To shed light on this phenomenon, here we compared the variance explained with CPM_CA_ for scans with left-right vs right-left PE direction including only scans from the first session of each subject. Interestingly, in the third column of Fig. 5, which corresponds to *t*-scores of correlations for left-right vs right-left PE direction, we observe antisymmetric patterns with respect to the anterior-posterior axis (see specifically the first three rows corresponding to axial slices). In other words, if for example a region in the left hemisphere was more prone to pulse-related oscillations for left-right PE direction compared to right-left PE direction, then the contralateral region in the right hemisphere was more prone to these oscillations for right-left PE direction. Moreover, we observe that the regions that exhibit this PE direction dependence were not necessarily the regions more susceptible to pulse-related oscillations.

**Fig. 5.**
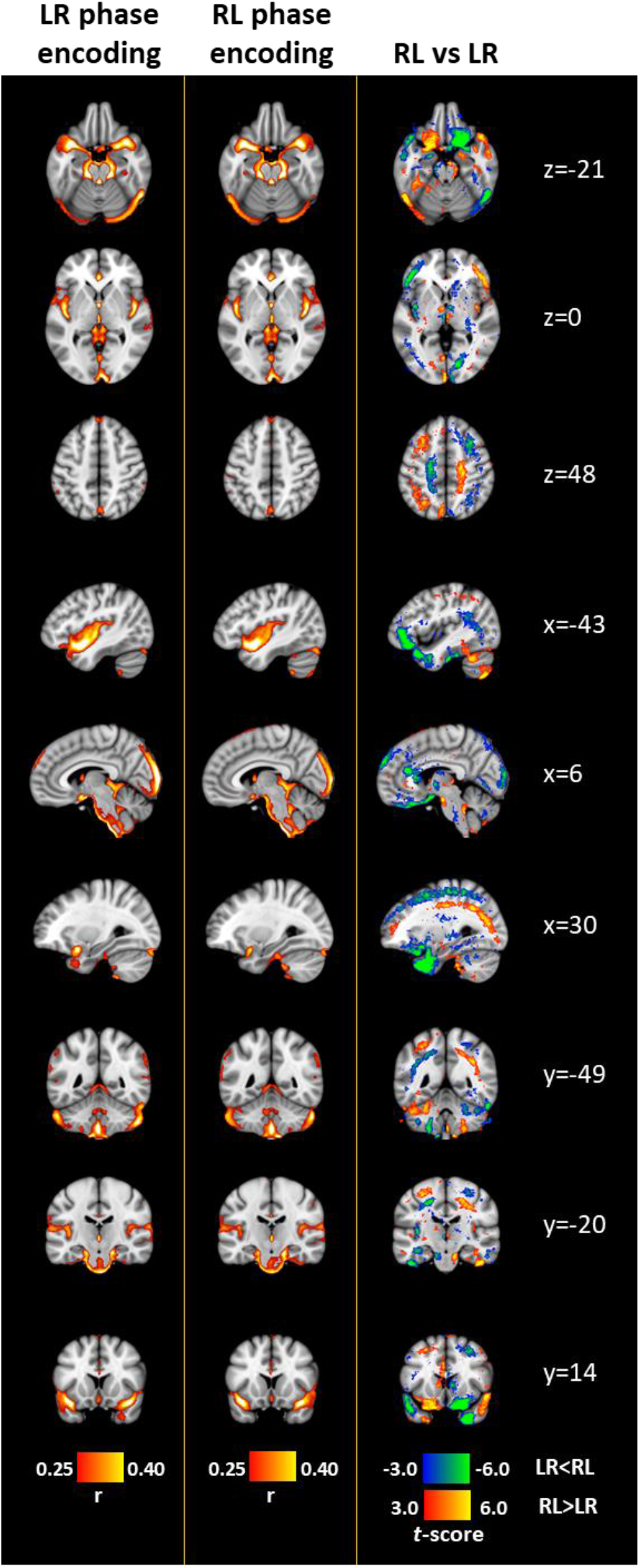
The effect of phase encoding (PE) direction on pulse-related fluctuations. The first and second columns show the variance explained using CPM_CA_ averaged across all subjects for scans with left-right and right-left PE direction, respectively (N=100; only scans from the first session were included here). The third column shows *t*-stat maps indicating the areas with significant differences between the two PE directions (*p*<0.01). The correlation threshold value was chosen arbitrarily so that only the regions more prone to cardiac pulsatility are presented (note that the cross-validation framework was omitted here, as we do not compare models but rather examine the effect of PE direction, hence the higher correlation values compared to Fig. 4). As can be seen from the axial slices, the regions with significant differences between the two PE directions were characterized by antisymmetric patterns with respect to the anterior-posterior axis. The statistical maps shown in this figure are available on: https://neurovault.org/collections/DHFETQTN/.

Furthermore, we investigated the regional variability of the CPWs. As described in Methods (Section 2.4.2), the beta parameters associated to the CPM nuisance regressors define the temporal waveforms of the impulse responses used in the CPM, referred to here as CPWs. Based on the beta parameters estimated with the GLM, we inspected the temporal evolution of the CPWs both at the individual (subject) and group (averaged across subjects considering only scans with the same PE direction from the first session) level. For this analysis, a 6^th^ order CPM_CA_ with a lag time of −0.9 s was considered, as the cross-validation analysis presented earlier showed that this choice of parameters yielded the best performance. At the individual level, we were not able to observe any clear pattern apart from discontinuities in adjacent slices that were likely due to time acquisition differences across slices. Note that we did not incorporate any slice-timing correction in our analysis as this is not a trivial task for the multi-band fMRI data examined here (Smith et al., 2013). However, at the group level (N=100), we were able to see smooth spatiotemporal patterns without discontinuities in adjacent slices. Videos showing the temporal evolution of CPWs in a mid-sagittal plane for left-right and right-left PE direction are available on repository https://doi.org/10.6084/m9.figshare.c.4946799 (Kassinopoulos and Mitsis, 2020b). In addition, Suppl. Fig. 4 shows the CPWs at five time points uniformly distributed within a cardiac cycle for the left-right PE direction. Similar dynamics were observed between the two PE directions. While some regions demonstrated an increase in the BOLD fMRI signal after the onset of a cardiac contraction (e.g. regions in the posterior cingulate cortex) other regions exhibited the opposite trend (e.g. third ventricle and regions in the anterior cingulate cortex). Interestingly, the temporal dynamics shown in the video revealed patterns that may relate to some fluid movement. Particularly, we observed two spatial layers with opposite temporal responses that peaked at about the middle of the cardiac cycle along the cerebral aqueduct that connects the 3^rd^ with the 4^th^ ventricle.

### 3.3 Systemic low-frequency oscillations and their relation to photoplethysmographic and global signal

Fig. 6 presents the results of the cross-correlation analysis that was used to investigate the relation between physiological variables (RV, HR and PPG-Amp) and the fMRI GS during resting conditions. Note that the cross-correlation technique can be used to estimate the impulse response for a single-input single-output system when the input signal is uncorrelated with the output noise (Lee and Schetzen, 1965; Marmarelis, 2004). However, in the present study, we used cross-correlation to assess the linear correlation patterns between the timeseries of interest, as well as the presence of a lag time between them. When examining the effects of SLFOs on the GS (presented later in this section), the estimation of the PRFs (impulse responses) that link the physiological variables RV, HR and PPG-Amp to the GS was done using the framework proposed in Kassinopoulos and Mitsis (2019), which employs basis expansion techniques and assumes a multiple-input single-output system.

**Fig. 6.**
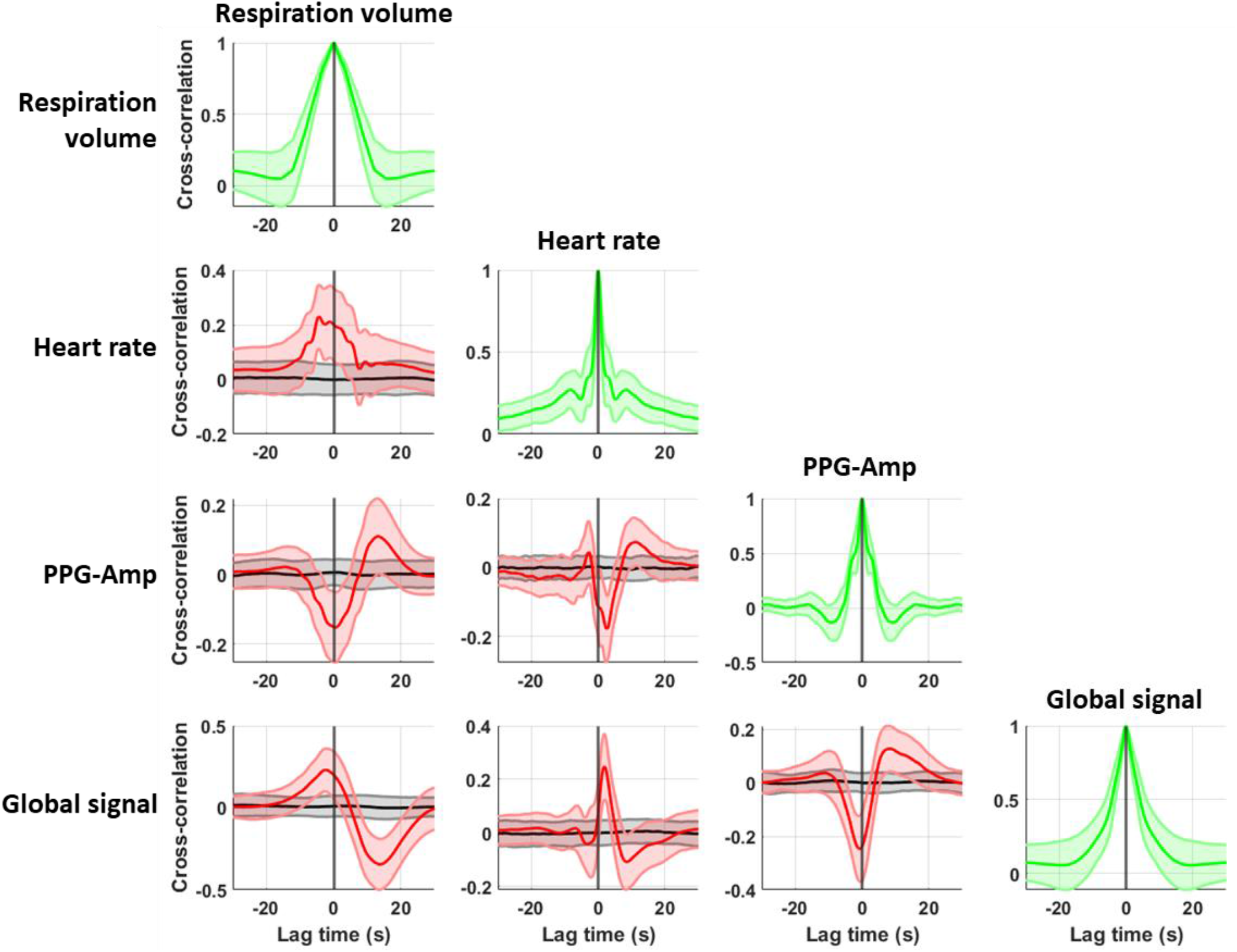
Cross-correlation between physiological variables and fMRI global signal (GS) averaged across subjects (N=100). The plot for row x and column y illustrates the auto-/cross-correlation of the corresponding signals. For instance, in the case of HR and GS, the fact that the maximum absolute cross-correlation occurs at +1.9 s and the correlation is positive indicate that HR is maximally correlated with the GS when it is shifted forward in time by 1.9 s. Shaded areas indicate the standard deviation across subjects and cross-correlation curves in red and black color correspond, respectively, to real and surrogate physiological variables (surrogate physiological variables were generated by phase randomization). Note that the diagonal plots correspond to autocorrelations. Overall, we observe that all physiological signals share covariance with each other as well as with the GS.

As can be seen in the diagonal of the diagram, all timeseries exhibited a somewhat monotonic decrease in autocorrelation for increasing (absolute) lag times with the autocorrelation approximating zero for lag times between 10 and 20 s. This trend is expected considering the sluggishness of the examined signals. Looking at the non-diagonal plots, we observe that, despite the low correlation values, all three physiological variables demonstrated interactions between them that were consistent across subjects. An increase in breathing activity, as quantified with RV, was accompanied by a concurrent increase in HR and decrease in PPG-Amp. However, PPG-Amp, apart from this decrease, exhibited also a positive peak about 13 s after the peak in RV. The cross-correlation between PPG-Amp and HR exhibited a similar bimodal form, albeit with different dynamics. Specifically, a negative peak was observed at about 2.5 s followed by a positive peak with a smaller amplitude at a lag time of 10 s.

In addition, GS was found to share covariance with all physiological variables. As expected, the cross-correlation functions between GS and RV or HR resemble, respectively, the CRF and RRF reported in our recent study (Kassinopoulos and Mitsis, 2019). As the main trend of the corresponding cross-correlations suggests, an increase in RV and HR leads to an increase in the GS followed by a negative undershoot. An opposite trend was observed in the cross-correlation between PPG-Amp and GS. Specifically, the cross-correlation between GS and PPG-Amp exhibited a strong negative peak at −1 s as well as a positive peak at 8 s.

The strong association observed between PPG-Amp and GS in the cross-correlation analysis (Fig. 6) raised the question whether considering PPG-Amp variations in addition to RV and HR variations, may provide additional information related to the effect of SLFOs on the GS. To address this question, we compared the standard approach for modeling SLFOs (i.e. considering only HR and RV) with the extended model that accounts also PPG-Amp variations, using a 3-fold cross-validation framework (for more information see Section 2.5). The cross-validation framework was necessary for this comparison due to the larger number of parameters in the extended model. Our results revealed a significant improvement in terms of variance explained in the GS when incorporating PPG-Amp variations. Specifically, the correlation between GS and predicted SLFOs exhibited a statistically significant increase from 0.64 (±0.10) to 0.65 (±0.10) when taking into account PPG-Amp variations (*p* < 10^−3^).

Suppl. Fig. 5a shows the weighted average PRFs derived from all subjects, specifically the cardiac (CRF), PPG-Amp (PARF) and respiration (RRF) response functions (see Section 2.5). Both CRF and RRF exhibited smooth bimodal curves with a positive peak followed by a negative peak and were in agreement in terms of dynamics with the PRFs reported in Kassinopoulos and Mitsis (2019a; Suppl. Fig. 6). In contrast, PARF demonstrated a sharp negative peak at 0.2 s followed by a slow positive overshoot.

The sharp negative peak in the PARF (Suppl. Fig. 5a) combined with the minimum peak observed at negative lag time in the cross-correlation between GS and PPG-Amp (Fig. 6) suggest that fluctuations in GS precede fluctuations in PPG-Amp. Because of this result, we also examined the extended SLFOs model with the PPG-Amp timeseries shifted back in time by 5 and 10 s. Below, we refer to the original, 5 s and 10 s shifted variants of PPG-Amp as PA_0_, PA_5_ and PA_10_, respectively. Using the cross-validation framework, PA_5_ was found to yield the highest mean correlation (0.66±0.09), which was significantly higher than the mean correlation (0.65) achieved with PA_0_ (*p* < 10^−4^). Fig. 7a shows the PRFs averaged across subjects when using the PPG-Amp shifted back in time by 5 s (PA_5_). As we can see, PARF exhibited a bimodal curve with a negative peak at 4.3 s, followed by a positive peak at 12.5 s. Due to the time shift in PPG-Amp, the negative peak of the PARF was smoother compared to the negative peak observed with the original PPG-Amp timeseries (Suppl. Fig. 5a). Moreover, the time shift in PPG-Amp did not have any effect on CRF and RRF. Similarly to our previous study (Kassinopoulos and Mitsis, 2019), we observed variability in the estimated curves across scans for all types of PRFs, with the CRF being the most consistent (the estimated PRFs from all scans examined here can be found on https://doi.org/10.6084/m9.figshare.c.4946799 (Kassinopoulos and Mitsis, 2020b)).

**Fig. 7.**
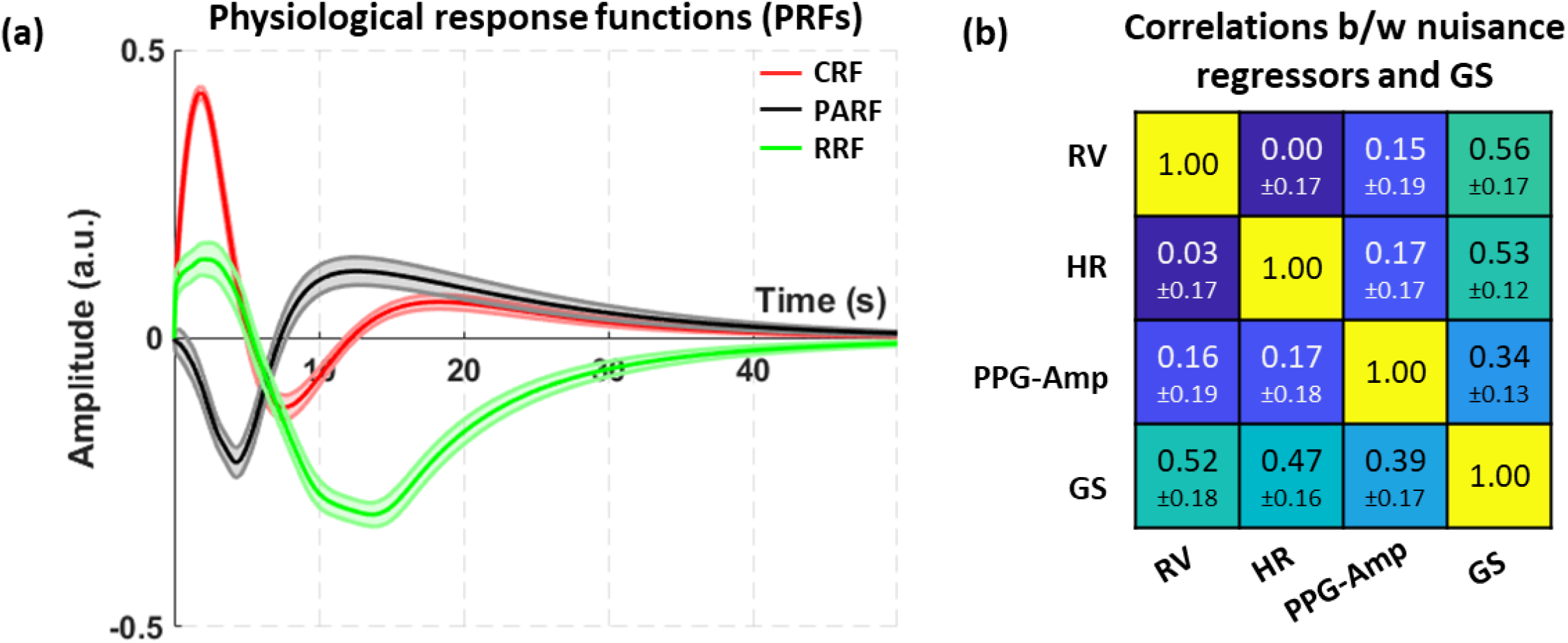
Estimated physiological response functions (PRFs) when considering the 5 s shifted PPG-Amp (PA_5_). **(a)** PRFs averaged across all subjects and scans using weighted averaging with the correlation between GS and the predicted output of the model (i.e. SLFOs) for each scan as the weighting coefficient. Shaded areas indicate the standard error across subjects. **(b)** Correlation between nuisance regressors (i.e. *X*_*RV*_, *X*_*HR*_and *X*_*PA*_) and GS, averaged across all scans, along with the standard deviation. The lower-diagonal elements correspond to correlations, whereas the upper-diagonal elements correspond to partial correlations. The partial correlations between pairs of the three nuisance regressors did not control for GS variations, as GS is not considered to affect the three associated physiological variables. Note that we do not compare models here, hence the cross-validation framework was omitted. CRF and RRF exhibited a positive peak at around 2 s followed by a negative peak at 8 s for CRF and 13-14 s for RRF. PARF was characterized by a negative peak at 4.3 s followed by a positive peak at 12.5 s. All PRFs demonstrated a slow decay that approximated zero at around 40 s. While the nuisance regressors demonstrated relatively low correlations between them ranging from 0.03 to 0.17, all of them exhibited high correlation with the GS (≥0.39). Similar observations were made for the partial correlations.

Fig. 7b shows the correlation averaged across all scans between the GS and the nuisance regressors extracted using the PRFs (*X*_*RV*_, *X*_*HR*_and *X*_*PA*_). As we can see, the mean correlation values between nuisance regressors were relatively low. However, for specific individual scans, we often observed pairs of nuisance regressors (*X*_*RV*_ vs *X*_*HR*_, *X*_*RV*_ vs *X*_*PA*_ and *X*_*HR*_vs *X*_*PA*_) that were highly correlated. Moreover, in Fig. 7b we see that all three nuisance regressors were strongly correlated with the GS, with *X*_*RV*_ exhibiting the highest mean correlation (0.52) and *X*_*PA*_ the lowest (0.39). While the mean correlation values between nuisance regressors were low, the cross-correlation analysis suggested strong interactions between the original physiological variables (Fig. 6). Due to this, we also computed partial correlation values to quantify the fraction of GS variance explained by each nuisance regressor, when removing the effect of the other two. Similar observations were made for the resulting partial correlation values (Fig. 7b). Fig. 8 shows the performance of the extended model for a scan where the nuisance regressors extracted from HR, PPG-Amp and RV were very similar and also explained a relatively large fraction of the low-frequency GS oscillations.

**Fig. 8.**
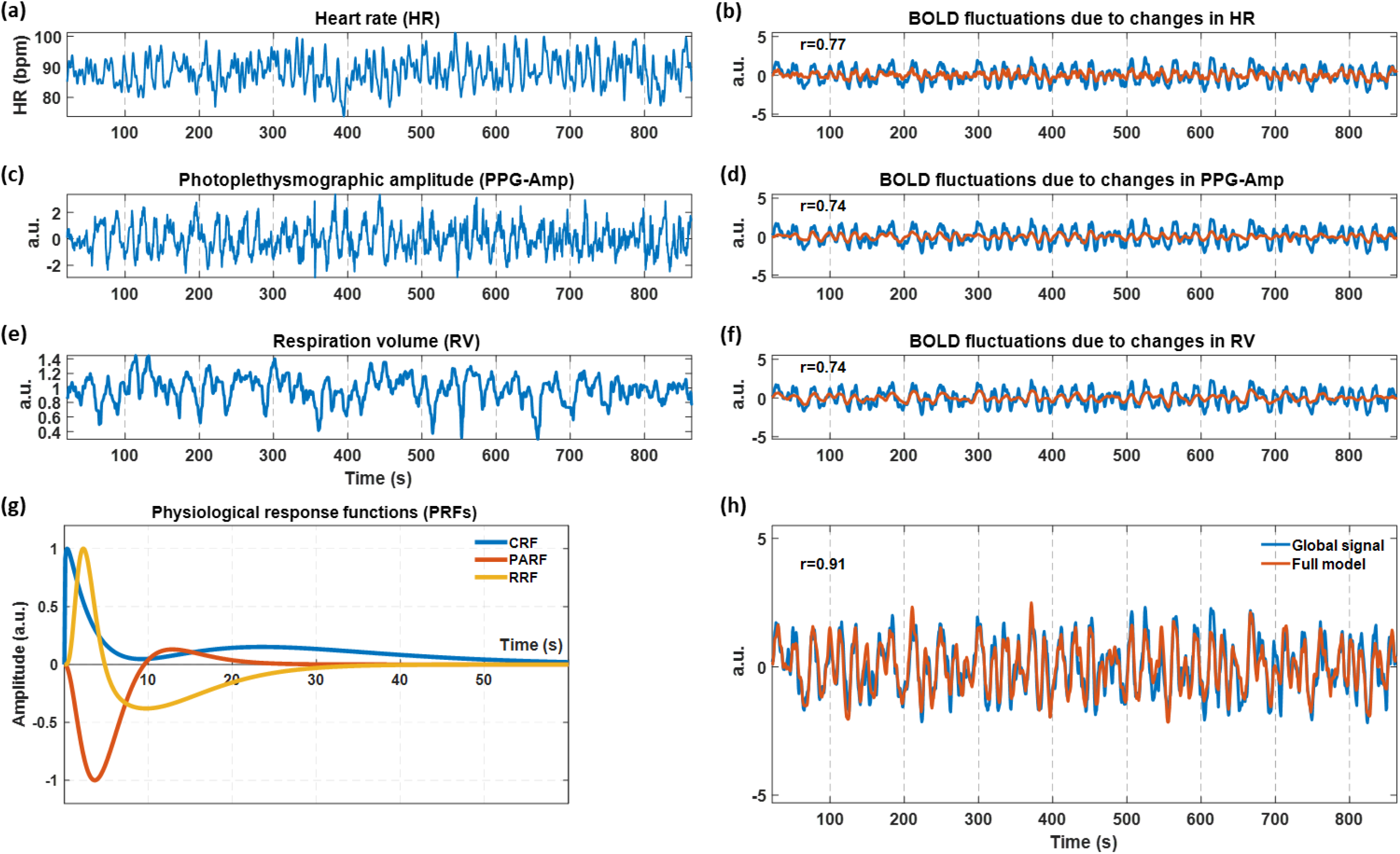
Demonstration of the estimated SLFOs in the GS for scan S103818-R2RL. **(a), (c)** and **(e)** show, respectively, the traces of HR, PPG-Amp and RV during the scan, whereas **(b), (d)** and **(f)** show the fit of the nuisance regressors extracted from the physiological variables (orange color) on the GS (blue color). **(g)** Scan-specific PRFs estimated using the framework proposed in Kassinopoulos and Mitsis (2019a). **(h)** Model fit of predicted output (i.e. SLFOs) on the fMRI GS. For visualization purposes, the HR in (a) was smoothed using a moving average filter of 3 s. For this particular scan, all three nuisance regressors obtained from HR, PPG-Amp and RV explained a large fraction of variance in the GS. The corresponding figures for all other scans can be found on https://doi.org/10.6084/m9.figshare.c.4946799 (Kassinopoulos and Mitsis, 2020b).

While the contribution of PPG-Amp variations to the GS was on average lower compared to the contribution of HR and RV variations, several subjects exhibited a stronger correlation between GS and PPG-Amp as compared to HR or RV. To shed light on this subject variability, we examined whether the partial correlation between PPG-Amp and GS depends on the body type and blood properties of the participants. Specifically, among the measures collected from participants in HCP, we considered the body weight, body mass index, height, systolic and diastolic pressure as well as hematocrit. In addition, we considered the HRV (i.e. standard deviation of HR) and standard deviation of RV as estimated from the physiological recordings. We chose these measures due to their strong correlation to hemodynamic properties such as blood viscosity and total peripheral resistance in the circulatory system. Among the eight tests performed, at a significance level of 0.05, only HRV and hematocrit were strongly associated to the partial correlation between GS and PPG-Amp (Fig. 9). Specifically, both exhibited a strong positive correlation to the partial correlation values between GS and PPG-Amp (*p*<0.001). Based on this observation, we also examined the relation between hematocrit and HRV. However, we did not find any significant correlation between them (*p*=0.67).

**Fig. 9.**
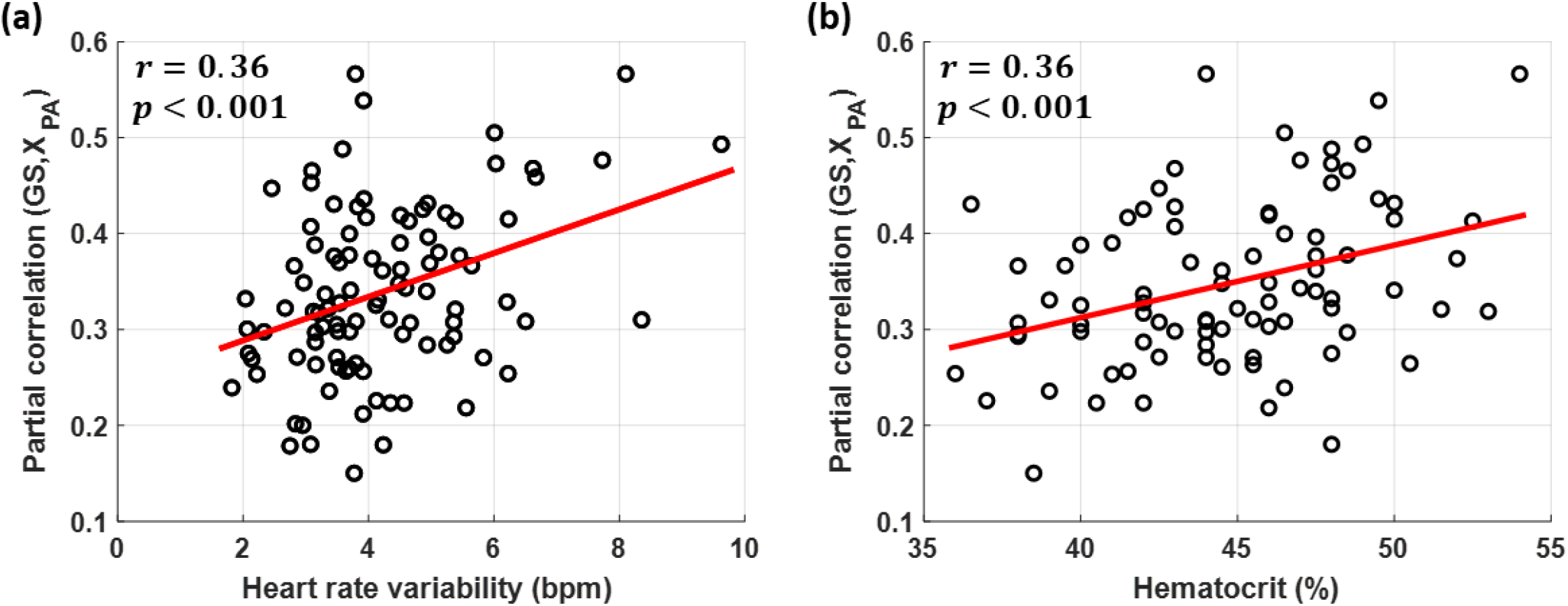
Scatterplots of the GS variance explained by PPG-Amp across subjects as a function of (a) HRV and (b) hematocrit. The hematocrit was measured for 87 out of 100 subjects. For subjects that had two measures of hematocrit, we used the mean value. The GS variance explained by PPG-Amp was calculated as the partial correlation between GS and the nuisance regressor *X*_*PA*_, averaged across the four scans of a subject. Similarly, HRV was averaged across scans for each subject. Interestingly, higher HRV and hematocrit values were associated with a larger fraction of GS variance explained by PPG-Amp.

Finally, consistent to previous studies, variations in HR and breathing pattern had a strong effect across widespread regions in the gray matter (Birn et al., 2006; Kassinopoulos and Mitsis, 2019; Shmueli et al., 2007). The same regions were found to be strongly associated with variations in PPG-Amp as well (Suppl. Fig. 7).

## 4. Discussion

In this study, we have revisited techniques commonly employed in the fMRI literature for modeling low-and high-frequency physiological-related fluctuations and proposed refinements and extensions. Furthermore, we sought to answer whether the low-frequency (∼0.1 Hz) amplitude modulations observed in the PPG recordings, referred to here as PPG-Amp, carry important information in the context of physiological noise modeling for either the low- or high-frequency physiological-related fluctuations.

### 4.1 High-frequency cardiac fluctuations

With regards to high-frequency oscillations induced by cardiac pulsatility, we considered Card-RETROICOR, which is a widely used technique proposed by Glover et al. (2000). Card-RETROICOR assumes that the pulse-related oscillations are described by a non-causal system as it requires knowledge of future input values (i.e. the timing of the following peak in PPG) to estimate the output at a specific timepoint. However, as the idea of considering a causal system that depends only on current and past input values is more physiologically plausible, we examined the feasibility of the convolution model CPM to capture pulse-related oscillations in fMRI. Importantly, a noise correction technique that considers a causal system can be directly implemented in real-time fMRI, such as in fMRI-based neurofeedback applications, addressing some of the limitations encountered with Card-RETROICOR with respect to the incremental update of the associated regressors (Misaki et al., 2015). Similar to Card-RETROICOR, CPM uses a Fourier series to construct a CPW for each voxel. The CPW in the proposed model plays essentially the role of an impulse response function consistent with the notion of hemodynamic or physiological response functions that has been adopted in the fMRI literature (Birn et al., 2008; Boynton et al., 2012; Chang et al., 2009). The input signal is defined as a train of pulses located at the timings of PPG peaks (Suppl. Fig. 1). Two variants of input signals were considered, one with all pulses having equal amplitude (value of one) and a second one where the amplitudes of the pulses matched the amplitudes of the PPG peaks. Note that the peaks observed in PPG are slightly delayed with respect to cardiac contractions due to pulse transit time effects (Allen, 2007). However, this time difference was accounted for when comparing Card-RETROICOR and CPM by considering the time shift that yielded the best performance for each model.

To compare the performance of the proposed CPM with Card-RETROICOR, we employed a 3-fold cross-validation framework. In addition, we sought to optimize the model order and determine whether incorporating some time shift in the PPG recordings can improve the performance in terms of BOLD variance explained in the voxel timeseries. Indeed, the time shifts of −0.4 s for Card-RETROICOR and −0.9 s for CPM were found to significantly increase the variance explained in the BOLD fMRI signal (Fig. 3a-c, Table 1). In the case of Card-RETROICOR, the time shift of −0.4 s is likely related to the fact that we considered the onset of an fMRI volume as the acquisition time for all slices in the volume. In the HCP fMRI data examined here, the TR which corresponds to the time interval between volumes was 0.72 s. However, as the slices of an fMRI volume were being acquired at different times during this time interval, we could consider that the effective time acquisition of a volume (i.e. the time that deviates the least from the acquired times of all slices) was the time indicated by the analog trigger shifted by half TR forward. If we had accounted for the aforementioned effective time acquisition, the optimal time shift for Card-RETROICOR would have been close to zero.

Jones et al. (2008) have suggested that slice-specific Card-RETROICOR regressors can explain more variance in the fMRI data compared to volume-wise regressors. However, the superiority of slice-specific regressors was demonstrated on fMRI data with a TR of 3 s and the improvement was marginally significant (*p* < 0.03). Therefore, it is still an open question whether the slice acquisition can have considerable implications for cardiac pulsatility modelling at fMRI datasets with shorter TR (e.g. 0.72 s). Even though it is of interest to examine the potential benefits of estimating regressors on a slice-wise basis, this was not feasible in the present study due to the non-trivial nature of slice acquisition in the pulse sequence used in the HCP (https://wiki.humanconnectome.org/display/PublicData/HCP+fMRI+slice-timing+acquisition+parameters). However, given that the cross-correlation curve obtained with CPM_CA_ for the fluctuations in the PPG timeseries has a broader maximum peak compared to RETROICOR (Fig. 2e), we speculate that CPM_CA_ would likely be less sensitive to slice timing accuracy compared to RETROICOR.

With respect to the half second difference in optimal time shift observed between CPM and Card-RETROICOR (Fig. 3a-c, Table 1), it may be related to the fact that the basis functions used in CPM were slightly altered compared to the basis functions used in Card-RETROICOR (specifically, the cosines were subtracted from one) so that the CPWs used in the convolution models, by design, started and ended at zero. Note that the two models showed a similar difference in optimal time shift when used to model the fluctuations in the PPG timeseries (Fig. 2e). With regards to optimal lag, we also observed that for both Card-RETROICOR and CPM, the cross-correlation curves were found to be monotonically decreasing around the maximum peak (Fig. 3a-c), even when considering individual voxels (Suppl. Fig. 3), suggesting that none of the models can account for time differences between physiological recordings and fMRI acquisition, unless the time delay is explicitly incorporated into the model.

Significant improvement in the variance explained was also observed when using higher order Fourier series compared to the 2^nd^ order typically used in the literature (Fig. 3d, Table 1). Specifically, for both Card-RETROICOR and CPM, the 6^th^ model order yielded the best goodness-of-fit values. Model orders higher than 6 demonstrated a small decreasing trend, suggesting that higher orders may result in overfitting and, thus, carry the risk of removing some signal of interest. While our results demonstrated the superiority of the 6^th^ model order compared to lower orders, we note that the optimal order may vary across datasets, and particularly across datasets with different pulse sequence parameters which determine the degrees of freedom in the data, such as TR and scan duration. Furthermore, even though we did not examine variability of optimal order across brain regions, we speculate that the optimal model order may be higher in regions prone to pulse-related artifacts, such as regions close to large arteries, ventricles and venous sinuses.

Among the three models examined in this work for pulse-related fMRI fluctuations, the proposed model CPM_CA_ exhibited the best performance in terms of variance explained (Fig. 3). As mentioned earlier, Card-RETROICOR and CPM_CA_ differ only in that the former assumes a CPW that is phase-locked to the cardiac cycle whereas the latter assumes a CPW of constant duration (Fig. 1). This suggests that the two models yield essentially the same output when the HR is relatively constant, and their outputs is likely to differ only in cases with strong fluctuations in HR. As hypothesized, the causal system CPM_CA_, which is more physiologically plausible than Card-RETROICOR, performed better than the latter for subjects with high HRV (above ∼5 bpm), whereas the performance of the two models was similar for subjects with low HRV (Fig. 3e). Based on this finding, we speculate that the benefits of using CPM over RETROICOR may be larger for studies that examine populations with high heart rate variability and task-based studies where the paradigm may have a strong influence on heart rate.

The rationale for examining CPM_VA_, which incorporates the low-frequency fluctuations in the PPG-Amp, was to examine whether the varying amplitude of cardiac pulses observed in the PPG (i.e. PPG-Amp) is also present in the BOLD fMRI signal. Compared to Card-RETROICOR and CPM_CA_, CPM_VA_ explained significantly less variance in the fMRI timeseries, suggesting that cardiac pulses measured in the brain with fMRI do not have the same amplitude as the pulses measured on the finger with PPG. We believe that the differences in amplitude variations observed between fMRI and PPG are not due to the different recording sites but rather to differences in the physical principles underlying each modality. PPG is based on near-infrared spectroscopy (NIRS), whereby a biological tissue is illuminated with near-infrared light from a laser diode and the light detected by a receiver in a nearby site is analyzed to provide information about the compounds present in the illuminated tissue (Delpy and Cope, 1997; Pellicer and Bravo, 2011; Scheeren et al., 2012). Based on the Beer-Lambert law, the attenuation of the light that is measured with PPG depends on the concentrations of oxyhemoglobin (O_2_Hb) and deoxyhemoglobin (dHb), as well as on their absorption coefficient for the wavelength of the incident light. Due to that O_2_Hb and dHb are characterized by different absorption spectra, it is very likely that a single-wavelength PPG shows different sensitivity to changes in O_2_Hb compared to dHb. Therefore, the variations observed in PPG-Amp may be the result of variations in the relative fractions of O_2_Hb and dHb which may not necessarily be accompanied by changes in the total hemoglobin (i.e. sum of O_2_Hb and dHb). On the other hand, while BOLD fMRI is considered to reflect changes in dHb, it is also very prone to motion artifacts. Therefore, the pulses in fMRI may originate mainly from arterial expansion and tissue movement due to the propagating blood pressure waves in the arteries rather than fluctuations in dHb. If this is indeed the case, then we would expect the pulse waveforms to be independent of the exact composition of blood (in other words, any fluid with similar viscosity to the blood would lead to similar fluctuations) and, therefore, independent of the relative changes in O_2_Hb and dHb. Another possible explanation for the differences in pulse amplitude between PPG and fMRI is that, while the PPG signal is linearly proportional to the levels of O_2_Hb and dHb, and subsequently to the total hemoglobin and blood volume, in fMRI the signal likely exhibits a non-linear relation to blood volume changes (Buxton et al., 1998; Friston et al., 2000). Therefore, variations in PPG-Amp due to changes in blood volume may be reflected differently in fMRI.

In a recent study, we demonstrated that cardiac pulsatility induces systematic biases in FC that to some extent depend on whether the PE direction is left-right or right-left (Xifra-Porxas et al., 2020). Here, we provide evidence that the regional sensitivity of fMRI data to cardiac pulsatility depends partly on PE direction (Fig. 5), which may explain why biases in FC due to cardiac pulsatility exhibit the same dependence. Note that dependence on PE direction was also reported for breathing motion fMRI artifacts by Raj et al. (2001). Based on simulations and experimental data, Raj et al. suggested that magnetic susceptibility variations, likely caused by lung expansion, induce variations in the static magnetic field within the brain being imaged (Raj et al., 2001, 2000). As a result, the spatial encoding during fMRI acquisition is unavoidably affected, leading to a shift of the reconstructed image in the PE direction as well as distortion of voxel timeseries with artifact waveforms that depend on both the phase of the breathing cycle and the location of each voxel with respect to the PE direction. In our study, the contribution of the pulse-related fluctuations varied across voxels, depending on their location with respect to the PE direction which may suggest that, similar to breathing motion, the mechanism by which pulsatility-induced vessel expansion gives rise to fluctuations in fMRI is partly through local variations in the static magnetic field. Note though that, as vessel expansion causes also fluid and tissue movement, spin-history effects are also thought to be another source of fluctuations (Caballero-Gaudes and Reynolds, 2017; Murphy et al., 2013).

The proposed model CPM_CA_ can be used to remove fMRI fluctuations due to cardiac pulsatility and facilitate the detection of neural-related activity. However, another potential application of this model is to visualize blood flow pulsatility in cerebral arteries as well as pulsatility-induced CSF movement. There is accumulating evidence that altered cardiac pulsatility in the brain is associated with neurodegenerative diseases such as Alzheimer’s disease (Harrison et al., 2018; Iliff et al., 2013; Mestre et al., 2018; Schley et al., 2006). As such, there is a growing interest in developing non-invasive techniques for measuring intracranial pulsatility. When we examined the CPWs (i.e. waveforms of the pulse-related fMRI fluctuations) extracted with CPM_CA_, we were able, at the group level, to observe patterns that resembled CSF movement to some extent, particularly in areas along the cerebral aqueduct (Suppl. Fig. 4, a video with the temporal dynamics is available on https://doi.org/10.6084/m9.figshare.c.4946799 (Kassinopoulos and Mitsis, 2020b)). This finding suggests that CPM_CA_, combined with a suitably designed fMRI pulse sequence, may be a potential tool for studying the pulsating brain.

### 4.2 Systemic low-frequency oscillations (SLFOs)

Using cross-correlation analysis, we showed that low-frequency GS fluctuations (∼0.1 Hz) were preceded by changes in physiological variables (RV, HR and PPG-Amp; Fig. 6). In addition, these three variables were, to some degree, associated to each other. When we considered convolution models to quantify their contributions to the GS, the latter was found to share unique variance with each of the three variables, with RV being the most influential factor and PPG-Amp the least influential (Fig. 7b).

The curves of the estimated RRFs and CRFs exhibited significant variability across scans. However, the main trend was similar to the trend reported in our earlier study (Suppl. Fig. 6; Kassinopoulos and Mitsis, 2019). Both CRF and RRF exhibited a positive peak at around 2 s followed by a negative peak at 8 s for CRF and 13-14 s for RRF. With regards to the shape of the CRF, as has been suggested in our previous study, the abrupt positive peak may reflect the increase in the blood flow that accompanies the HR increase, whereas the negative peak a few seconds later may reflect a regulatory feedback mechanism, potentially mediated by a decrease in stroke volume, that aims to bring the blood flow back to its baseline, despite the changes in HR. On the other hand, the early positive peak in RRF may indicate that an increase in breathing activity, either due to increase in breathing rate or breathing volume, leads to an abrupt increase in O_2_Hb, which results in an increase in the BOLD signal. However, increased breathing activity leads also to a decline in arterial CO_2_. As CO_2_ is a strong vasodilator, its decrease leads to a somewhat delayed vasoconstriction which, in turn, reduces the blood flow and also the BOLD signal.

Inspired by the earlier studies of Birn et al. (2008) and Chang et al. (2009), here we introduced an impulse response function, termed photoplethysmographic amplitude response function (PARF), that relates PPG-Amp variations to changes in GS. To achieve the best fit on the GS, the PPG-Amp was shifted backward by 5 s. A negative time-shift for RV and HR was not considered when modeling the SLFOs in the GS, as the cardiac and breathing activity are processes thought to drive GS fluctuations. On the other hand, the PPG signal collected from the participant’s finger, as with the BOLD signal, can be seen as a hemodynamic signal whose fluctuations are controlled, among others, by the cardiac and breathing activity. Therefore, the negative time-shift needed for the PPG signal may suggest that the blood pumped by the heart to the aorta arrives earlier at the brain vasculature than to the finger. Note that even though the cross-correlation between GS and PPG-Amp analysis yielded a peak at −1 s, when examining the SLFOs model with the PPG-Amp as one of the three inputs, shifting the PPG-Amp back in time by 1 s led to a small, albeit not significant increase in the GS variance explained. Due to this, we also examined the SLFOs model with the PPG-Amp shifted backward by longer time intervals and found that a time-shift of 5 s results in a small, yet significant increase in the variance explained compared to the original PPG-Amp (*p* < 10^−4^), and also yields a smoother PARF curve (Fig. 7a vs Suppl. Fig. 5a).

While the improved performance observed with the 5 s time-shift may be related to differences in the blood arrival time from the heart to the brain vs the finger, it may also be related to the use of the double gamma function for modeling the PARF. The double gamma function enforces a PRF curve to start at zero amplitude which, while desirable for the RRF and CRF, may not be a good choice for the PARF. Transcranial Doppler (TCD) studies have shown that the impulse response that describes the effects of arterial blood pressure on cerebral blood flow is characterized by a large amplitude at time zero (Kostoglou et al., 2014; Mitsis et al., 2004). Therefore, considering the possible association of PPG-Amp and GS with changes in arterial blood pressure and cerebral blood flow, respectively, a PARF allowed to start at a non-negative value may be more appropriate for modelling the associated BOLD responses. In addition, given the potential instantaneous effects of PPG-Amp on the GS, the negative time-shift may improve the goodness-of-fit on the GS by compensating somehow for the non-ideal choice of basis functions.

The shape of the estimated PARF averaged across all subjects exhibited a somewhat opposite trend compared to CRF and RRF. Specifically, it exhibited a negative peak at 4.3 s followed by a positive peak at 12.5 s (Fig. 7a). The shape of PARF cannot be easily interpreted, as PPG-Amp reflects several processes. Consistent with previous studies (Özbay et al., 2019, 2018), PPG-Amp was found to exhibit an almost instantaneous drop when HR increased (Fig. 6), which may be associated to decreased stroke volume. Moreover, PPG-Amp was found in our data to be reduced during inhalation, a phenomenon well-documented in the literature that has been attributed to changes in intrathoracic pressure resulting also in a reduced stroke volume (Meredith et al., 2012). In addition, as PPG is sensitive to changes in HbO and dHb, it also captures slower effects of cardiac and breathing activity related to changes in blood oxygenation and volume. While PARF may be lacking a clear physiological interpretation, the cross-validation analysis conducted in the present study revealed that the inclusion of PPG-Amp convolved with PARF in the model of SLFOs improved the goodness-of-fit on the GS, compared to considering only HR and RV. Specifically, in the cross-validation analysis the mean correlation increased from 0.65 to 0.67, which was found to be statistically significant (*p*<10^−7^), while when the model was trained and tested on the same dataset the mean correlation increased from 0.73 to 0.76 (*p*<10^−26^).

The contribution of PPG-Amp on the GS exhibited variability across subjects, with partial correlation values between PPG-Amp and GS ranging between 0.15 and 0.57. Subjects with higher HRV were characterized by a stronger relationship between GS and PPG-Amp (Fig. 9a; *p*<0.001), supporting the notion that PPG-Amp explains variance on the GS induced partly by HR changes. Furthermore, we observed that the higher was the hematocrit of a subject, the larger was the contribution of PPG-Amp on the GS (Fig. 9b; *p*<0.001). The hematocrit, which is defined as the proportion of red blood cells in the blood, is considered as one of the factors determining the amplitude of the PPG signal even though its exact effect is still not very clear (Fine, 2014; Jubran, 2015; Ochoa and Ohara, 1980). The strong relationship between hematocrit and contribution of PPG-Amp to the GS can be explained as follows: higher levels of hematocrit lead to a stronger relative effect of HbO and dHb on the PPG signal, compared to other compounds. As a result, the PPG becomes more sensitive to changes in oxygenation, which in turn leads to an increased variance explained in the GS using the amplitude of the PPG pulses for subjects with high hematocrit levels. Note that a positive linear relationship has been previously reported between the amplitude of task-induced BOLD responses and the hematocrit (Gustard et al., 2003; Levin et al., 2001), which would suggest that the mean or standard deviation of the GS may differ between subjects with different hematocrit values. However, when we examined the mean and standard deviation in the GS, we did not find any strong association with hematocrit values (results not shown).

The capability of PPG-Amp to remove SLFOs from fMRI data was first demonstrated by Van Houdt et al. (2010), who showed that removing these fluctuations facilitates the detection of the epileptogenic zone in patients with epilepsy. Van Houdt et al. reported a low negative correlation between RVT (a measure of breathing activity similar to RV used here) and PPG-Amp (−0.08±0.09), even though similar brain regions were influenced by the two variables. While we also found similarly low values in the cross-correlation analysis, the trends were consistent across subjects (Fig. 6). Importantly, consistent with Van Houdt et al. (2010), we found that the regions associated to variations in PPG-Amp (widespread regions across GM; Suppl. Fig. 7) were similar to the regions associated to variations in HR and RV (Kassinopoulos and Mitsis, 2019). This may not be surprising, as we expect physiological processes to affect areas close to the vasculature, and particularly close to draining veins, such as in the occipital cortex, where the concentration of dHb varies the most. That said, the three variables seem to consist of shared but also unique variance, hence the increased variance explained when considering all variables in the analysis.

A main difference between the analysis employed here and the analysis in Van Houdt et al. (2010) is that we accounted for the dynamics for the effects of physiological processes on fMRI data using convolution models and basis expansion techniques. By doing so, we ensured the plausibility of the estimated PRFs while also keeping the number of free model parameters low. In contrast, Van Houdt et al. (2010) considered 10 lagged versions of the PPG-Amp in the GLM to account for the underlying dynamics. This multi-lagged approach reduces degrees of freedom in the data, particularly in the frequency range where we expect neuronal-related activity (Bright et al., 2017) and, therefore, is prone to removing signal of interest. On a related note, in an earlier study we found that there was no benefit when allowing variability in the shape of the PRFs across voxels (Kassinopoulos and Mitsis, 2019). Therefore, as proposed by Falahpour et al. (2013), we chose to estimate subject-specific PRFs only from the GS which has high signal-to-noise ratio and therefore it is less likely to overfit the data compared to when estimating the PRFs on a voxel-wise manner.

A strong association of PPG-Amp with the GS was shown in recent sleep studies by means of cross-correlation analysis (Özbay et al., 2019, 2018). Specifically, Özbay et al. (2019, 2018) provided evidence that an increase in PPG-Amp is followed by an increase in the GS with a lag of a few seconds. In the present study, we found that the lagged increase in the GS followed by an increased in PPG-Amp occurs also during resting conditions. In addition, we also found a strong negative (cross-)correlation between PPG-Amp and GS for lag times between −4 s and 2 s that was not observed during sleep. While we find the aforementioned difference between the two studies puzzling, this difference may be somehow explained by the fact that the PPG signal is driven by several physiological processes (e.g. HRV) which in turn exhibit different trends across stages of sleep and wakefulness (Elsenbruch et al., 1999). As the role of PPG-Amp in fMRI has been somewhat neglected in the literature, further research is needed to shed light on the mechanisms that may determine the relation between PPG-Amp and GS as well as how this relation may vary between stages of sleep and wakefulness.

In the present study, we considered the pulsatile component of PPG as well as the variations in its amplitude to refine previously proposed models of physiological noise in fMRI. This component, often termed alternating current (AC) component in the NIRS literature, is commonly used to obtain measurements of HR and breathing rate (Charlton et al., 2018), and has been shown to explain fMRI variance related to quasi-periodic cardiac (∼1.0 Hz) and breathing (∼0.25 Hz) artifacts (Verstynen and Deshpande, 2011) as well as low-frequency oscillations (<0.1 Hz) induced by variations in breathing patterns (Van Houdt et al., 2010). However, another important component of the PPG that has not been well-investigated in the fMRI literature is the direct current (DC) baseline of the PPG signal at frequencies below 0.2 Hz. As a matter of fact, this DC component is typically removed from MRI pulse oximeters using a high-pass filter to facilitate the visualization of cardiac pulses (this is also likely the case with the PPG signals in HCP as they are lacking a low-frequency content). Pulse oximeters considering the full spectrum of the PPG signal that also illuminate at more than one wavelengths can provide additional physiological variables compared to single-wavelength PPG signals, such as variations in relative changes of HbO and dHb, as well as oxygen saturation (Delpy and Cope, 1997). Due to these properties, the DC baseline in PPG is of great importance in clinical cardiovascular monitoring (Jubran, 2015). Note also that the two-wavelength PPG signal is the fundamental signal exploited in functional NIRS (fNIRS) for the study of brain activity (Tachtsidis and Scholkmann, 2016).

Although fMRI studies do not typically consider the low-frequency (<0.2 Hz) fluctuations in PPG, its potential in physiological noise correction has been demonstrated by Tong and Frederick (2010). Specifically, Tong and Frederick (2010) demonstrated that the low-frequency fluctuations in HbO and dHb, obtained from peripheral NIRS, explained a significant variance in BOLD fMRI, while in a subsequent study they provided evidence that HbO and dHb explained a significantly higher variance compared to nuisance regressors related to HR and breathing patterns (Hocke et al., 2016). As the authors stated, this result may not be surprising as HbO and dHb measured with NIRS are physiological variables directly related to the BOLD signal. While the levels of HbO and dHb measured from the finger are affected by cardiac and breathing activity, they are also affected by other processes such as fluctuations in blood pressure and activity of autonomic nervous system that are non-trivial in terms of measurement, especially in the MR environment. Therefore, measurements from peripheral NIRS can in principle enable us to account for several factors in addition to HR and breathing pattern variations. With regards to our results, it is very likely that the unique variance explained by PPG-Amp in the GS could also be captured with the low-frequency fluctuations in NIRS. However, in case that only the PPG is available, the PPG-Amp could be considered in addition to the effects of HR and breathing pattern.

## 5. Conclusion

We examined noise correction techniques that utilize the PPG signal in order to account for low- and high-frequency physiological fluctuations in BOLD fMRI. The CPM_CA_ model was proposed as a physiologically plausible refinement of RETROICOR. CPM_CA_ employs a convolution framework in a similar manner to the use of hemodynamic response functions for modeling neural-induced BOLD responses. As initially hypothesized, CPM_CA_ performed equally well with Card-RETROICOR for subjects with relatively stable HR and outperformed the latter for subjects with high variability in HR. Moreover, we examined the potential implications of the variations in PPG-Amp (i.e. pulse amplitude observed in the PPG) on both low- and high-frequency physiological fluctuations. Our results suggest that the variations in PPG-Amp do not covary with the amplitude in the fMRI pulse-related fluctuations, but they explain a significant amount of variance in SLFOs present in the GS, in addition to variance explained by fluctuations in HR and breathing patterns. Given that the PPG-Amp reflects variations in the relative levels of oxyhemoglobin (O_2_Hb) and deoxyhemoglobin (dHb), our findings support the notion that pulse-related artifacts arise due to vessel expansion and associated motion artifacts in the nearby tissue compartments rather than to changes on blood oxygenation, whereas SLFOs are driven by physiological processes through changes in blood oxygenation. Overall, the pulsatile component of the PPG signal was found to explain a large fraction of variance in fMRI related to both low- and high-frequency physiological fluctuations. Scripts for the techniques proposed here are available on git repository https://github.com/mkassinopoulos/Noise_modeling_based_on_PPG.

## Acknowledgments

This work was supported by the Natural Sciences and Engineering Research Council of Canada (Discovery Grant 34362 awarded to GDM), the Fonds de la Recherche du Quebec - Nature et Technologies (FRQNT; Team Grant PR191780-2016 awarded to GDM) and the Canada First Research Excellence Fund (awarded to McGill University for the Healthy Brains for Healthy Lives initiative). MK acknowledges funding from Québec Bio-imaging Network (QBIN). Data were provided by the Human Connectome Project, WU-Minn Consortium (Principal Investigators: David Van Essen and Kamil Ugurbil; 1U54MH091657) funded by the 16 NIH Institutes and Centers that support the NIH Blueprint for Neuroscience Research; and by the McDonnell Center for Systems Neuroscience at Washington University. The authors would like to thank Alba Xifra-Porxas for her assistance in the selection of subjects from the HCP database with good quality physiological recordings.

## Supplementary Material

**Suppl. Fig. 1.**
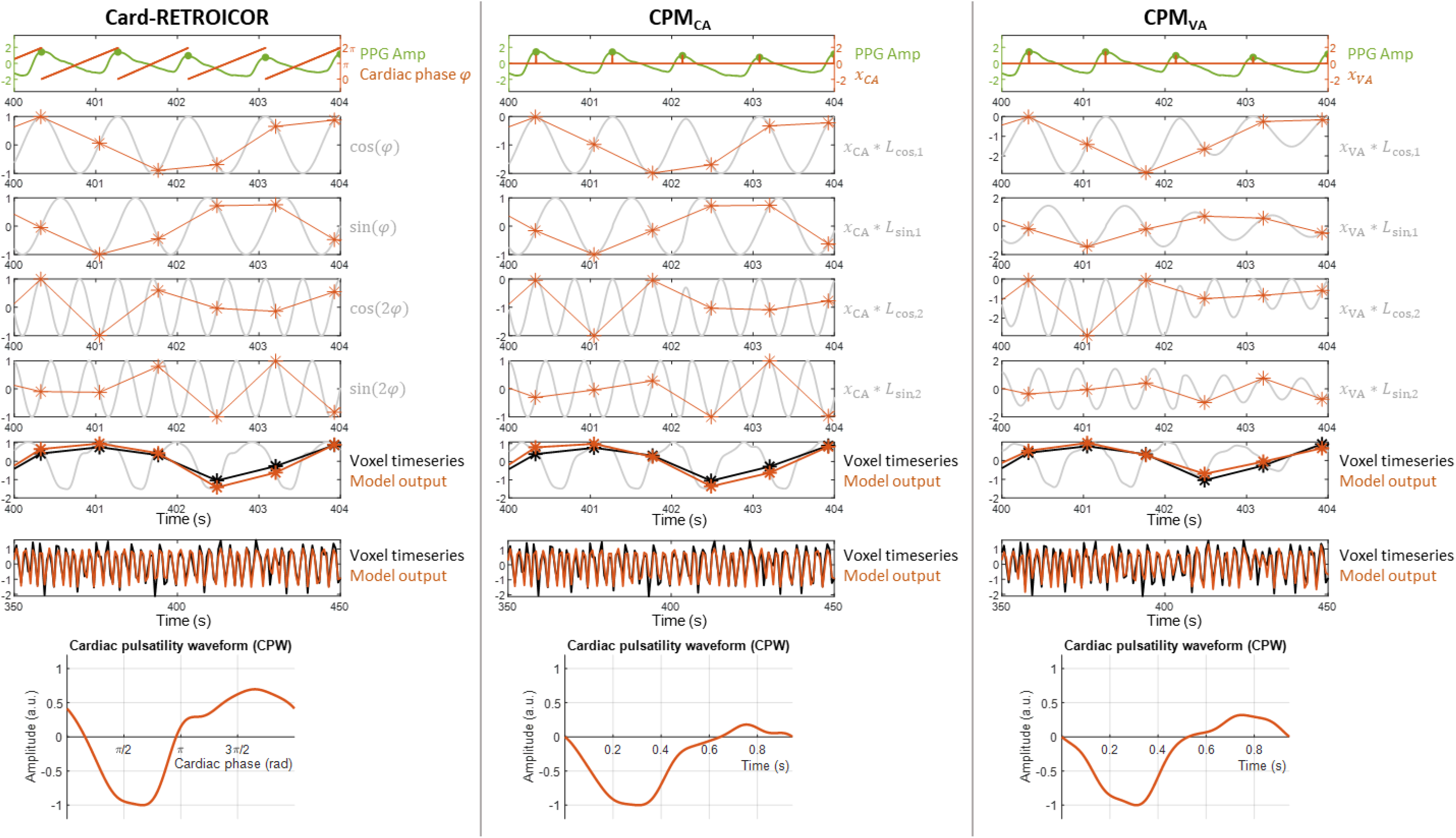
Demonstration of the main steps for modeling pulse-related fMRI fluctuations using Card-RETROICOR, CPM_CA_, and CPM_VA_ (left, middle and right column, respectively). For Card-RETROICOR, the cardiac phase *φ* is first computed based on the times of the PPG peaks (1^st^ row; *φ* is considered to increase linearly from 0 to 2π between adjacent peaks). Subsequently, a basis set of cosine and sine functions is created based on the cardiac phase and resampled at the times when fMRI volumes are acquired (2^nd^ to 5^th^ row; the stars (*) indicate the times of the fMRI volumes; for visualization purposes only the first four sinusoids are shown here). Finally, a linear combination of the resampled sinusoids is determined through linear regression that best explains the voxel timeseries (6^th^ and 7^th^ row; the timeseries correspond to a voxel nearby the brainstem from scan S515014_Rest1LR that was found to be strongly driven by cardiac pulses). Note that as the repetition time TR (0.72s) in HCP fMRI data does not satisfy the Nyquist criterion, the high-frequency (∼1.0 Hz) cardiac oscillations observed in PPG are reflected on fMRI timeseries as oscillations at lower frequencies (∼0.4 Hz), a phenomenon known as *aliasing*. The linear combination of sinusoids used to model pulse-related fluctuations can be applied to the Fourier basis functions to construct the cardiac pulsatility waveform (8_th_ row). For CPM_CA_, a train of impulses located at the times of the PPG peaks is first constructed and used as an input signal *x*_*CA*_(*t*)in the model (1^st^ row). This input signal is subsequently convolved with Fourier basis functions and the output of this convolution is resampled at the times of the fMRI volumes (2^nd^ to 5^th^ row). Finally, similar to RETROICOR, a linear combination of the resampled sinusoids is determined through linear regression that best explains the voxel timeseries (6^th^ and 7^th^ row). CPM_VA_ only differs from CPM_CA_ in that the input signal *x*_*VA*_ consists of a train of pulses with amplitudes equal to the amplitudes of the PPG peaks.

**Suppl. Fig. 2.**
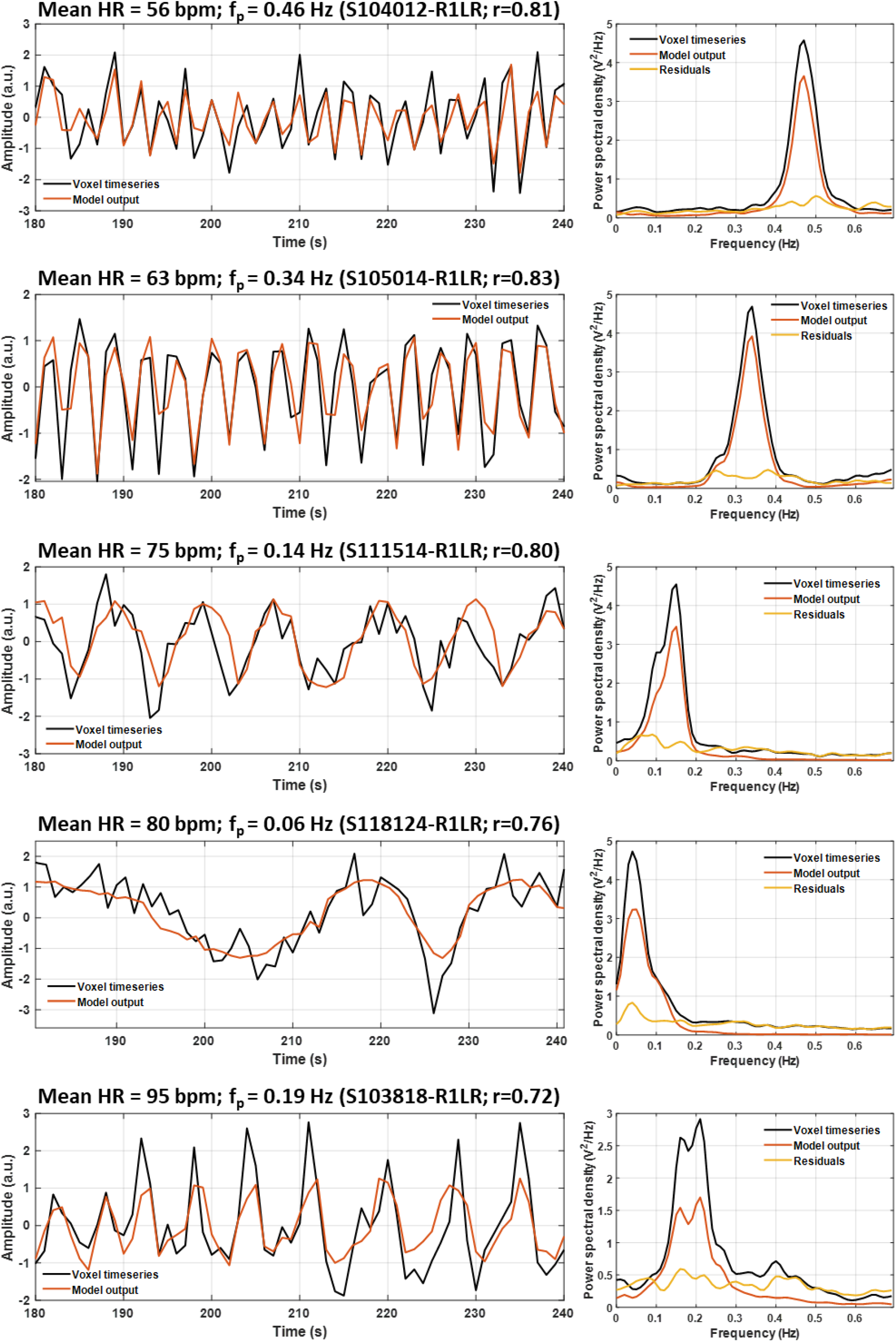
fMRI voxel timeseries, model output (left column) and their corresponding power spectral density (PSD; right column) for five subjects with different mean heart rate (HR). For each subject, the voxel with the strongest pulse-related fluctuations is presented (i.e. the voxel that demonstrated the highest correlation for the fit obtained with CPM_CA_). Due to the value of TR (0.72 s), the cardiac oscillations in the fMRI timeseries are aliased to low frequencies with the aliased (perceived) frequency f_p_ varying across individuals depending on their HR (subjects are sorted from low to high HR to better demonstrate the dependence of the aliased frequency to the HR).

**Suppl. Fig. 3.**
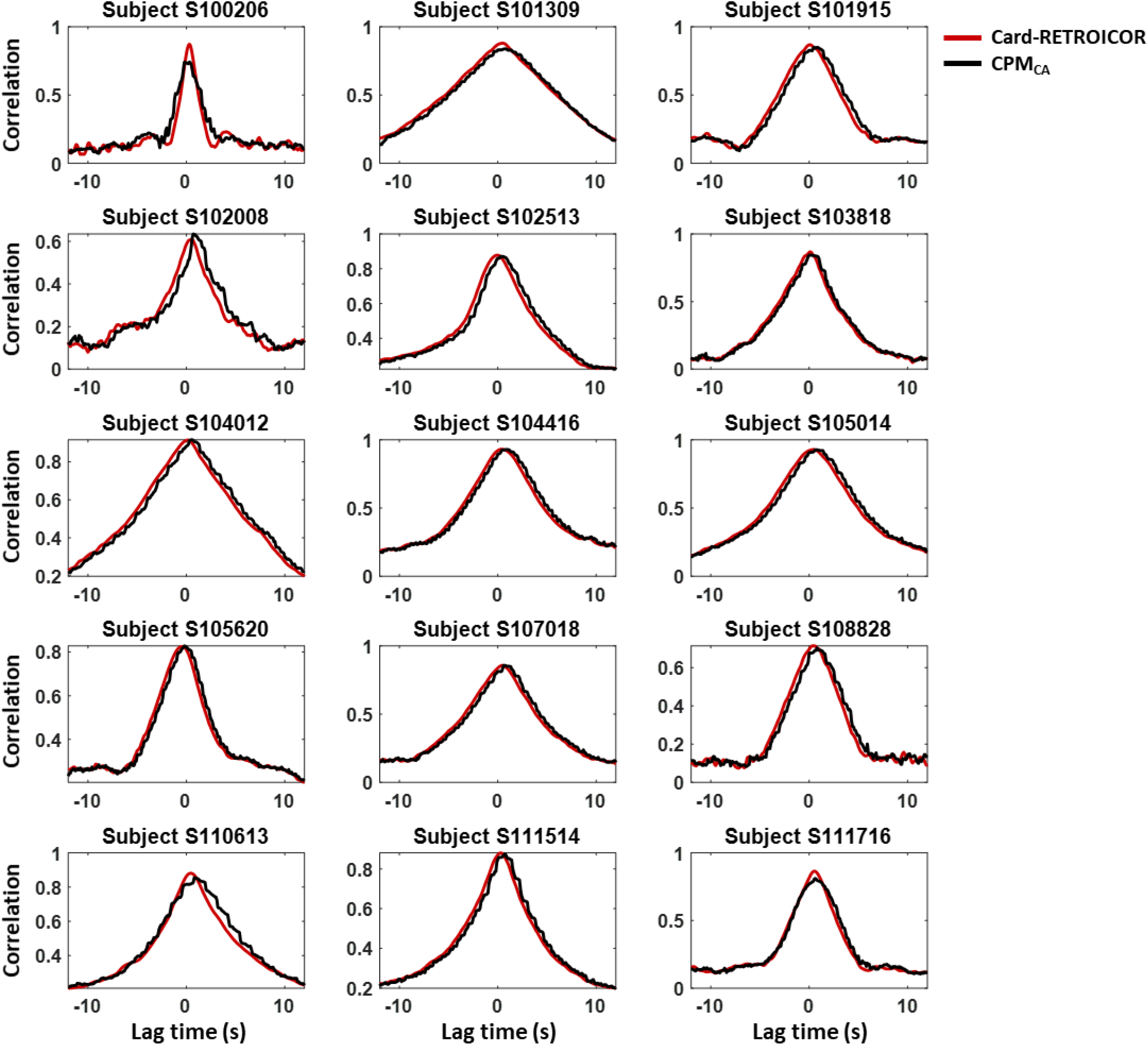
Performance of RETROICOR and CPM_CA_ in individual voxels for varying lag times. The cross-correlation for the voxel with the highest correlation is shown for 15 representative subjects. Both models demonstrate a monotonically decreasing cross-correlation with maximum peaks at around zero lag.

**Suppl. Fig. 4.**
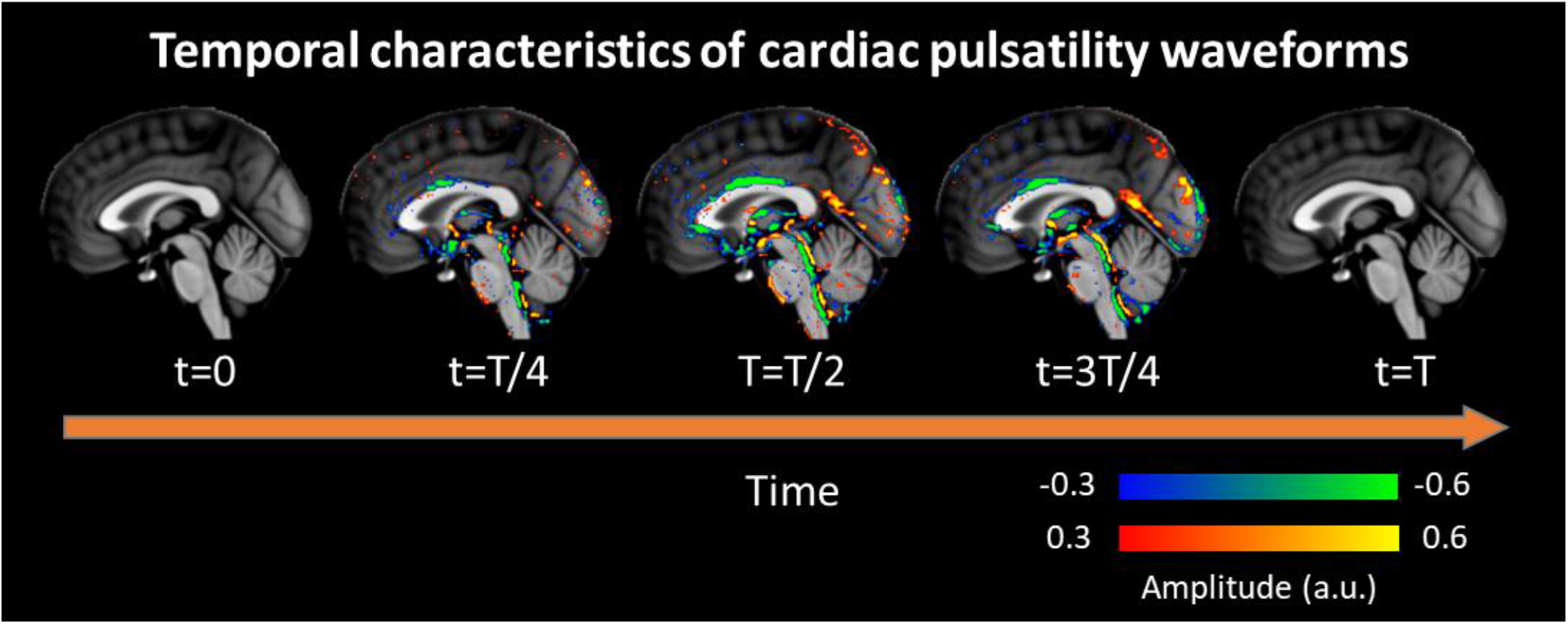
Cardiac pulsatility waveforms (CPWs) averaged across all subjects (N=100; only scans with left-right PE direction from the first session were included). Several areas such as the 3^rd^ ventricle and the cerebral aqueduct, as well as areas in the anterior and posterior cingulate cortex exhibited similar CPWs across subjects. A video presenting these waveforms induced by cardiac pulsatility in BOLD fMRI can be found on https://doi.org/10.6084/m9.figshare.c.4946799 (Kassinopoulos and Mitsis, 2020b). Intriguingly, the CPWs reconstructed with CPM_CA_ revealed different dynamics across regions which may be related to fluid movements.

**Suppl. Fig. 5.**
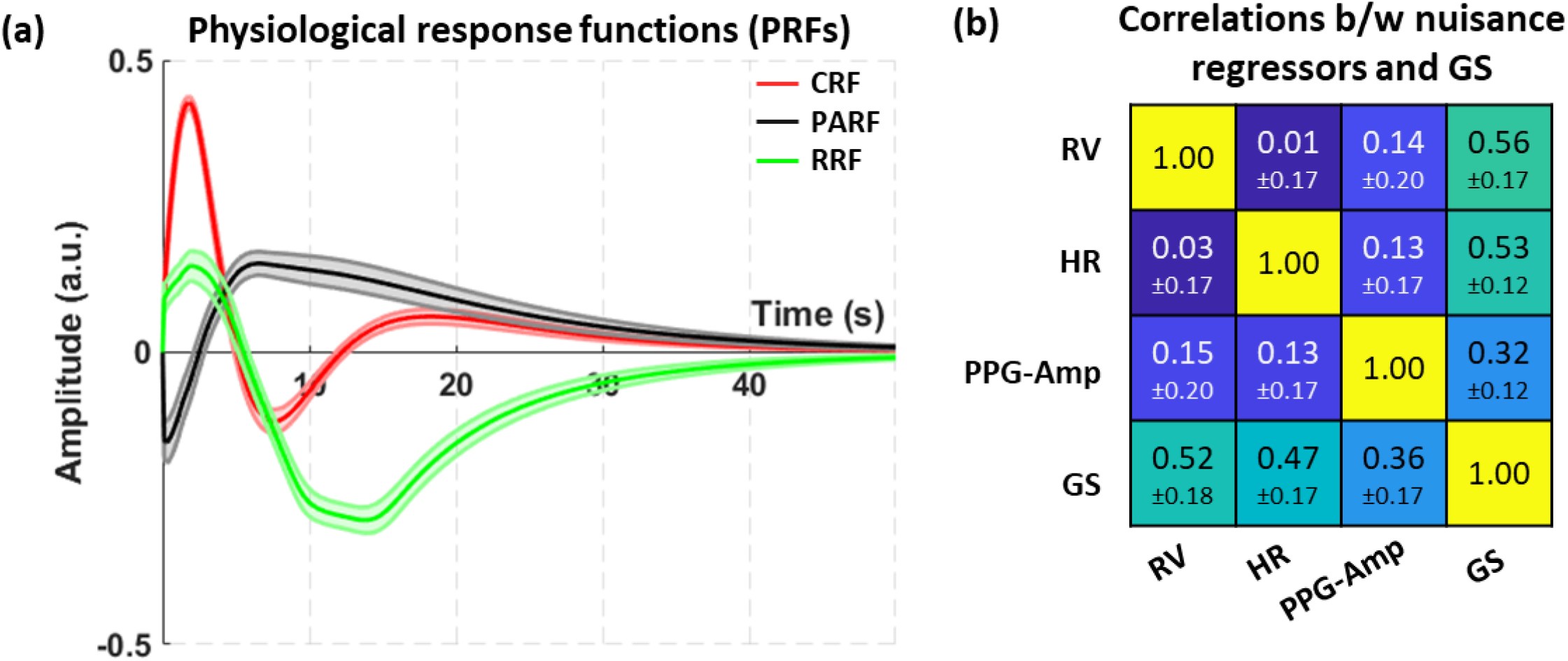
Estimated physiological response functions (PRFs) when considering the original, non-shifted PPG-Amp timeseries (PA_0_). (a) PRFs averaged across all subjects and scans using weighted averaging, with the correlation between GS and the predicted output of the model (i.e. SLFOs) for each scan as a weighting coefficient. Shaded areas indicate the standard error across subjects. (b) Correlation between nuisance regressors (*X*_*RV*_, *X*_*HR*_and *X*_*PA*_) and GS, averaged across all scans, along with the standard deviation. The lower-diagonal elements correspond to correlations, whereas the upper-diagonal elements correspond to partial correlations. The partial correlations between pairs of the three nuisance regressors did not control for GS variations, as it was assumed that GS did not affect the three associated physiological variables. Note that we do not compare models here, hence the cross-validation framework was omitted. CRF and RRF exhibited a positive peak at around 2 s, followed by a negative peak at 8 s for CRF and 13-14 s for RRF. PARF was characterized by a negative peak at 0.2 s followed by a positive peak at 6.5 s. All PRFs demonstrated a slow decay that approached zero at around 40 s. While the nuisance regressors yielded relatively low correlation values (between 0.03 and 0.15), all were strongly correlated with the GS (≥0.36). Similar observations were made for the partial correlations.

**Suppl. Fig. 6.**
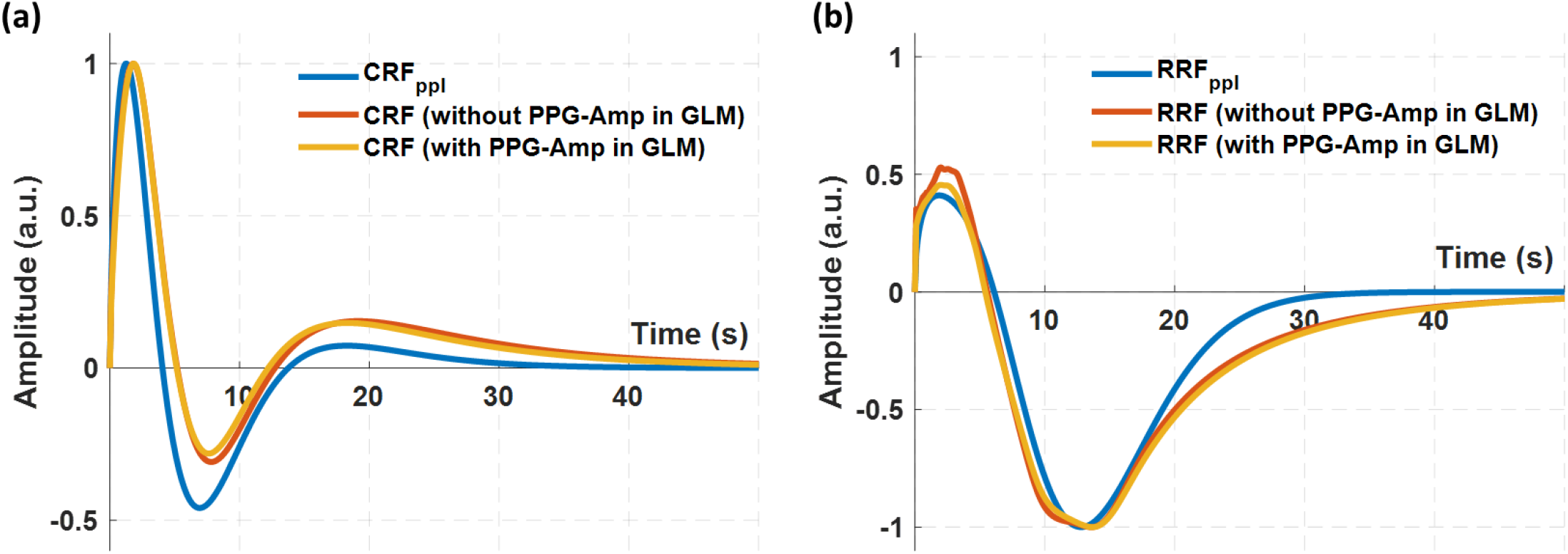
Weighted average of (a) cardiac and (b) respiration response functions (CRF and RRF, respectively) with and without considering the shifted PPG-Amp timeseries (PA_5_) in the general linear model (GLM). For both CRF and RRF, the inclusion of the PPG-Amp in the GLM did not have a considerable effect on the curves. The weighted average CRF and RRF curves exhibited only slight differences with the population-specific curves CRF_ppl_ and RRF_ppl_ (blue color) reported in Kassinopoulos & Mitsis (2019), presumably due to the different way in which they were obtained. Specifically, in the present study, scan-specific PRF curves were averaged across all subjects whereas in Kassinopoulos & Mitsis (2019) single population-specific CRF_ppl_ and RRF_ppl_ curves were estimated. Note also that a different respiratory measure was used as input variable in the two studies (respiratory flow vs respiration volume).

**Suppl. Fig. 7.**
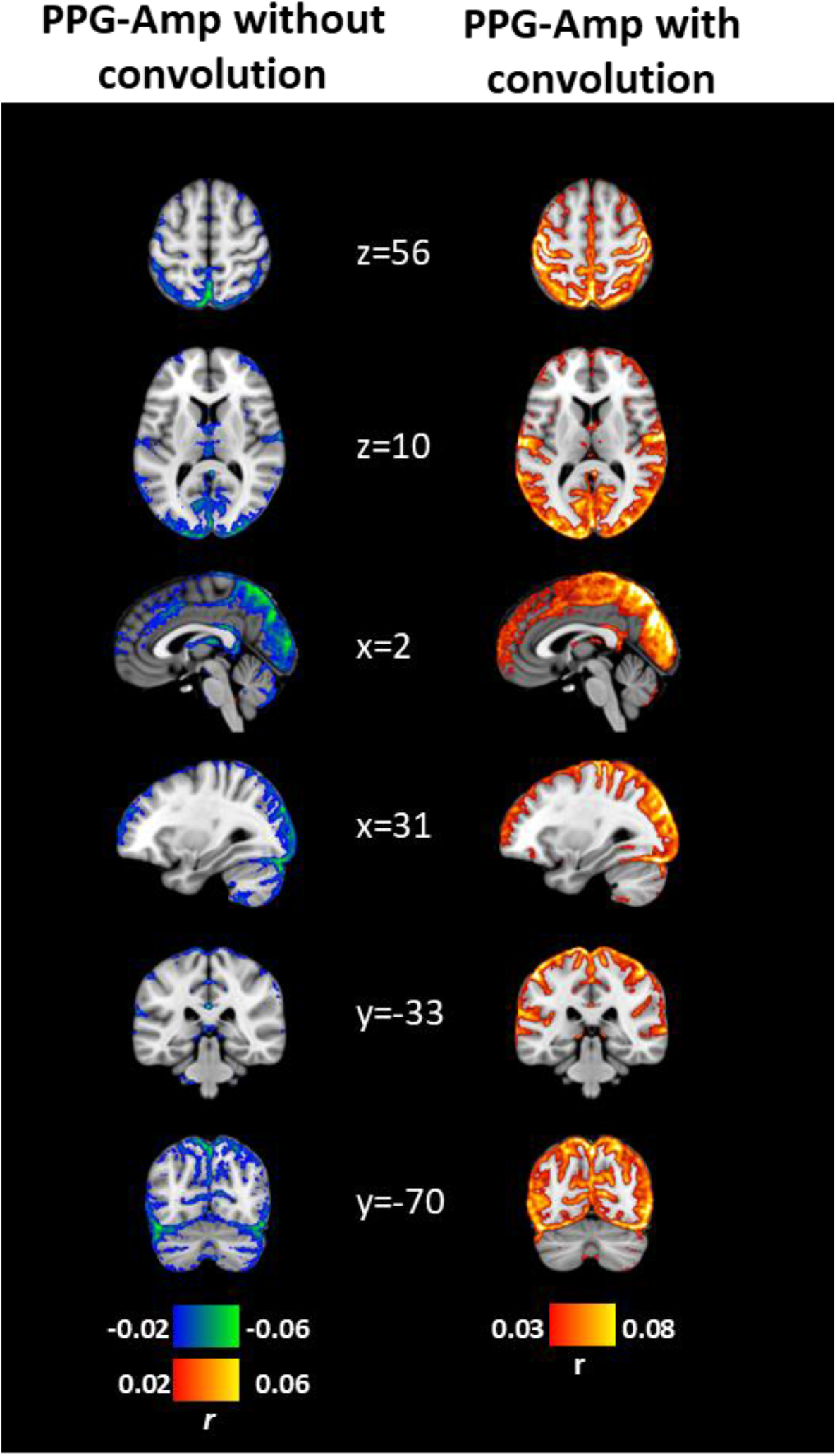
Contribution of PPG-Amp variations in fMRI averaged across all subjects. (1^st^ column) Correlation maps related to the original PPG-Amp variable. (2^nd^ column) Correlation maps related to the nuisance regressor *X*_*PA*_ extracted by shifting the PPG-Amp back in time by 5 s and convolving it with the PARF. The correlation maps were estimated on fMRI data corrected for head and breathing motion as well as cardiac pulsatility (6^th^ order of CPM). We observe that both the original PPG-Amp variable and its associated nuisance regressor explained variance in widespread regions across GM. However, as we can see, in the case of the nuisance regressor, the correlation values were significantly higher.

## Notes

### Competing Interest Statement

The authors have declared no competing interest.

